# Nanoscale Structural Mapping of Protein Aggregates in Live Cells Modeling Huntington’s Disease

**DOI:** 10.1101/2023.10.09.561223

**Authors:** Zhongyue Guo, Giulio Chiesa, Jiaze Yin, Adam Sanford, Stefan Meier, Ahmad S. Khalil, Ji-Xin Cheng

## Abstract

Protein aggregation, in the form of amyloid fibrils, is intimately correlated with many neurodegenerative diseases. Despite recent advances in structural biology, it remains challenging to acquire structural information of proteins in live cells. Tagging with fluorescent proteins, like green fluorescent protein (GFP), is routinely used for protein visualization. Yet, this method alone cannot provide detailed structural information on the protein system of interest, and tagging proteins has the potential to perturb native structure and function. Here, by fluorescence-detected as well as label-free scattering-based mid-infrared photothermal (MIP) microscopy, we demonstrate nanoscale mapping of secondary structure of protein aggregates in a yeast model of Huntington’s disease. We first used GFP as a highly sensitive photothermal reporter to validate β-sheet enrichment in huntingtin (htt) protein aggregates. We then obtained label-free structural maps of protein aggregates. Our data showed that the fluorescent protein tag indeed perturbed the secondary structure of the aggregate, evident by a spectral shift. Live cell MIP spectroscopy further revealed the fine spatial distribution of structurally distinct components in protein aggregates, featuring a 246-nm diameter core highly enriched in β-sheet surrounded by a ɑ-helix-rich shell. Interestingly, this structural partition exists only in presence of the [*RNQ*^+^] prion, a prion that acts to facilitate the formation of other amyloid prions. Indeed, when htt is induced to aggregate in the absence of this prion ([*rnq*^-^] state), it forms non-toxic amyloid aggregates exclusively. These results showcase the potential of MIP for unveiling detailed and subtle structural information on protein systems in live cells.

**Significance:** Protein aggregation is a hallmark of neurodegenerative diseases, such as Huntington’s Disease. Understanding the nature of neurotoxic aggregates could lead to better therapeutic approaches. The limited progress in this direction is partly due to the lack of tools for extracting structural information in the physiological context of the aggregates. Here, we report a photothermally detected mid-infrared micro-spectroscopy technique able to dissect the secondary structure of aggregates of the huntingtin protein in live cells. We describe for the first time a nanoscale partition of secondary structures between β-rich core and ɑ-rich shell of the aggregates. This work demonstrates the potential of mid-infrared photothermal microscopy for structural and functional mapping of proteins in live cells.

## Introduction

Neurodegenerative diseases (NDs) are a common and growing cause of mortality, morbidity and cognitive impairment worldwide, with no effective treatments or diagnostic biomarkers available. The incidence of most neurodegenerative diseases increases dramatically with advancing age (1), leading to rising financial, societal and emotional costs in our global ageing population. One common pathogenic mechanism underlying a wide variety of neurodegenerative diseases involves the accumulation of insoluble protein aggregates in the form of amyloid fibrils (2, 3). For instance, accumulation of amyloid-β and tau are associated with Alzheimer’s disease, whereas aggregates of ɑ-synuclein are found in Parkinson’s disease patients (4, 5). The same protein could also assemble into different structures, a phenomenon called structural polymorphism, and lead to different disease outcomes (5, 6). The intimate mechanism of amyloid fibrils formation remains unclear: protein monomers may first assemble into oligomers, which further assemble into fibrils by a nucleation-growth mechanism (4). Liquid-liquid phase separation (LLPS) has been invoked as an alternative mechanism for pathological protein aggregation (7, 8).

Despite the long standing interest in protein aggregation, structural studies of protein aggregates in live cells remain challenging (9). Conventional spectroscopic approaches (10, 11) and biophysical characterizations (12–14) have been extensively applied to fibrils *in vitro*. Recent breakthrough in solid-state nuclear magnetic resonance spectroscopy (ssNMR) (15, 16) and cryo-electron microscopy (cryo-EM) (17–19) led to determination of the atomic structure of amyloid fibrils. However, the pathological relevance of these isolated fibrils studied *in vitro* remains unclear, since it does not take in account the complex environment in which aggregates form and elongate.

Intracellular protein aggregates are widely studied by using fluorescent dyes (20, 21), or in fusion with a fluorescent protein, further aided by advanced microscopy (22, 23) and genetic engineering (9, 24–26). However, fluorescence itself could not provide structural information. Additionally, potential oligomerization and aggregation mediated by the tagged fluorescent protein may further complicate the study (27). Recently, cryo-electron tomography (cryo-ET) has been used to visualize the 3D structure of polyQ aggregates *in situ* in yeasts (28, 29). However, due to the complicated preparation of frozen hydrated and vitrified cell samples, this technique only allows one snapshot in time.

Recently developed mid-infrared photothermal (MIP) microscopy (30–33), also called optical photothermal infrared (O-PTIR) microscopy, opens a new way to structural studies of protein aggregates in live cells. MIP is a vibrational spectroscopic imaging technique, where a visible probe beam detects the photothermal effect induced by the mid-infrared (mid-IR) absorption. MIP simultaneously offers high spatial resolution from the visible probe, and rich chemical and structural information from the mid-infrared pump (30–33). MIP imaging of subcellular structures in live cells has been successfully demonstrated in cancer cells and neural cells (30, 34–37). Moreover, the quantitative spectroscopic nature of MIP makes it particularly suitable to study the structure of protein aggregates. The protein amide I band arising mainly from the C=O stretching vibration of the protein backbone correlates predictably to the protein secondary structure and it has long been used in conventional Fourier transform infrared (FTIR) analysis *in vitro* (38–41). Using a commercial mIRage O-PTIR microscope, Oxana Klementieva and coworkers mapped β-sheet aggregation by intensity ratio of the 1630 cm^-1^ peak over the 1650 cm^-1^ peak in primary neurons cultured with synthetic Aβ(1–42) and showed structural polymorphic amyloid-β aggregates in AD transgenic neurons (42). More recently, Jian Zhao et al. developed a fluorescence-guided bond-selective intensity diffraction tomography and achieved 3D visualization of intracellular tau fibrils by β-sheet structures in human cells seeded with extracted tau fibril fractions (43). Craig Prater et al. combined wide-field epi-fluorescence imaging with O-PTIR to perform IR spectroscopic analysis with the guidance of fluorescent amyloid tracers or amyloid-specific antibodies (44). However, these studies were limited to micrometer-scale aggregates in fixed cells or cryo-dried tissues. The full potential of MIP for nanometer-scale structural analysis in live cells remains untapped.

Here, using a live yeast model of Huntington’s Disease, we demonstrate MIP structural mapping of nanoscale protein aggregates. We recognize that water absorption through the bending vibration mode constitutes a critical challenge in the amide I window. To alleviate this challenge, instead of using co-propagation as in earlier work (42, 44), where the probe beam is weakly focused by a reflective objective, we built a counter-propagating MIP system equipped with a high numerical aperture refractive objective for the visible probe beam to enable photothermal detection of nanosized protein aggregates. Moreover, the very tightly focused probe beam does not sense the sizable thermal lens formed by IR excitation of bulk water. This advantage minimizes the background from medium and enabled selective detection of locally concentrated protein aggregates in live cells. In addition, epi-fluorescence excited by the same probe beam is added to the system for fluorescence-guided MIP spectral analysis of target proteins in live cells. Furthermore, we have developed a wide-field fluorescence-detected MIP setup and harnessed GFP as a highly sensitive photothermal reporter for high-throughput mapping of host protein secondary structure. Combining these strategies, we successfully dissected the secondary structure of nanoscale aggregates and described for the first time a partition of β-sheet rich core and ɑ-helix rich shell within the aggregation complex. We started with fluorescently labeled protein Huntingtin in yeasts. Harnessing the thermal sensitivity of fluorophores including GFP (45, 46), we first performed wide-field fluorescence-detected MIP and confirmed β-sheet enrichment in aggregates in a high throughput manner. We then used fluorescence as a guidance to identify protein aggregates in live cells and performed MIP spectroscopic analysis. We found significant spectral changes in protein aggregates (i.e. within fluorescent foci) that correspond to enrichment in β-sheet. Based on these spectral features, we achieved label-free identification of protein aggregates and detected a further red shift in spectra, indicating perturbations caused by the fluorescent protein tag. Next, multispectral MIP imaging revealed a spatial distribution of structurally distinct components within the protein aggregation complex, with a core highly enriched in β-sheet and a shell dominant by ɑ-helix. Finally, we tested how the yeast prion [*RNQ^+^*] affects the structure of the aggregates, by bypassing its nucleation activity with a recently developed optogenetic approach (47). We observed that in a [*rnq^-^*] background, htt forms smaller, non-toxic and exclusively β-sheet aggregates that do not assemble into larger round aggregates as in the [*RNQ^+^*] background. These data provide a structural explanation of the causes of toxicity in htt aggregates in the model organism *Saccharomyces. cerevisiae*. Altogether, our work demonstrates the potential of MIP spectroscopy for structural dissection of protein aggregation processes in live cells.

## Results

### Principle of MIP analysis of protein secondary structure in live cells

The concept of MIP secondary structural analysis of protein aggregates in live cells is illustrated in **Fig. 1*A***. The protein aggregates in their native environment are studied by live cell MIP imaging, using yeast as the eukaryotic model of protein aggregation. The mid-infrared light provides rich chemical and structural information while the tightly focused visible probe light provides high spatial resolution that enables imaging of small protein aggregates inside live yeast cells. To acquire secondary structure information of protein aggregates, we analyzed MIP signals at the wavenumbers corresponding to the protein amide I band. The amide I band vibration originates mainly from the C=O stretching in the peptide bonds of the protein backbone, and it is directly associated to the backbone conformation, making it most sensitive to protein secondary structure (39, 41). Second derivative analysis and deconvolution of the amide I band reveal component peaks, which can be assigned to different secondary structures (38). MIP spectrum is faithful to the conventional IR absorption spectrum and allows quantitative measurements for structural study, as the MIP signal is linearly proportional to the number density of molecules (30). Therefore, quantifiable secondary structural analysis of protein aggregates in live yeast cells could be achieved by MIP.

**Fig.1.**
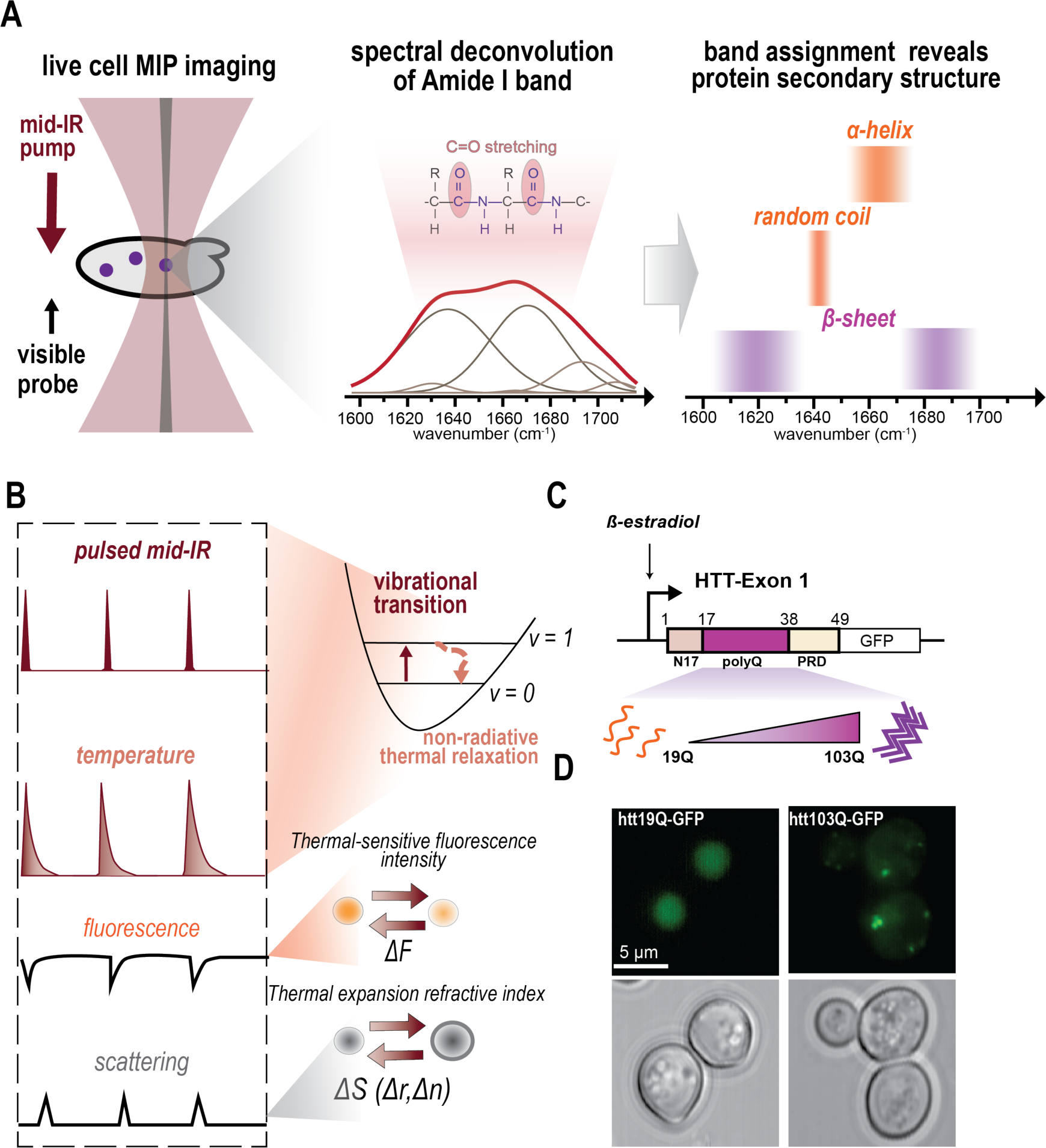
Secondary structural analysis of protein aggregates in live cells using a counter-propagating MIP system. (*A*) Concept of secondary structural analysis of protein aggregates in live cells. MIP imaging of live cells provides high spatial resolution to focus on small protein aggregates while mid-IR absorption in the amide I band reveals protein secondary structures after deconvolution and component band assignment. (*B*) Representation of MIP detection: mid-IR pulses induce both fluorescence and scattering intensity changes. (*C*) β-estradiol inducible model for Huntington’s Disease and protein aggregation in yeast. The sequence representation of HTT- Exon 1 is based on the exon 1 of htt protein (*homo sapiens*, UniProt P42858). PolyQ length predictably affects aggregation propensity. Low polyQ length (19Q) leads to soluble protein while high polyQ length (103Q) leads to protein aggregates. *(D*) Example of yeasts expressing htt19Q- GFP show diffuse fluorescence while yeasts expressing htt103Q-GFP show bright fluorescent foci indicating protein aggregates. Top: GFP fluorescence images. Bottom: corresponding transmission images. Scale bar: 5 µm.

The contrast mechanism of MIP imaging is explained in **Fig. 1*B***. Two laser beams are focused to the same spot: one mid-infrared and one visible. The selective absorption of mid-infrared light by molecules causes a local temperature change. This photothermal effect origins from the vibrational transition after absorbing mid-infrared light, and the subsequent nonradiative thermal relaxation (33). The sample is heated up locally by the short mid-infrared pulses, and gradually cools down through thermal diffusion. As fluorescence intensity decreases with increasing temperature, such that the photothermal effect can be detected by fluorescence (45). Alternatively, this temperature change leads to thermal expansion and a change in refractive index, detectable by measuring the differential in scattering (30). Hence, detecting mid-infrared absorption by visible light greatly improves spatial resolution while providing rich chemical information.

### A eukaryotic model of protein aggregation

To develop and benchmark the application of MIP spectroscopy for the exploration of secondary structure of protein aggregates, we leveraged a well-established model for protein aggregation in *Saccharomyces cerevisiae*, also called budding yeast, that recapitulates certain aspects of pathology in neurodegenerative diseases (**Fig. 1*C***) owing to a remarkably high degree of conservation in pathways related to protein quality control between yeast and higher eukaryotes. Since onset of neurodegenerative diseases is intimately associated with degeneration of cell proteostasis, yeast is a particularly effective model system for this type of disorders (48). Also, yeast enables introducing genetic manipulations simply, rapidly and inexpensively (49). Huntington’s Disease (HD) is a hereditary disease, caused by an expanded CAG trinucleotide repeat in the exon 1 of HTT gene, coding for huntingtin (htt) protein. The codon expansion leads to an abnormally long polyglutamine (polyQ) sequence in the mutant htt protein (50). There is a well-established correlation between polyQ length of the htt protein, its propensity to form insoluble aggregates and age of onset of the pathology (51–53): the larger the polyQ length, the more aggregation prone is the protein bearing it. This property can be recapitulated also in *S. cerevisiae*, establishing yeast as a model for HD (49).

In our system, exon 1 of protein huntingtin (htt) is conditionally overexpressed in a strain of *S. cerevisiae*. Overexpression of htt is known to cause aggregate formation, recognizable as puncta or foci by Thioflavin-S staining (or fluorescent foci, when the protein is tagged with a fluorophore) (54, 55). The protein fragment formed by exon 1 retains both the intrinsically disordered 17 N-terminal amino acids (N17) and the proline-rich region directly C-terminal to the polyQ. We considered a suite of mutants with polyQ length ranging from 19Q to 159Q. We then fused htt to a fluorescent protein (htt-FP). To regulate htt expression, we used a recently developed β-estradiol induction system (**Fig. 1*C***) (56). This orthogonal inducer for htt expression reduces confounding effects, such as metabolic changes due to change of carbon source and acute cytotoxicity (57). We confirmed that htt-FP expressed in this setting would cause toxicity to cells, by monitoring growth curves of cells expressing either htt19Q-FP (non-aggregation prone) or htt103Q-FP (highly aggregation-prone). Indeed, htt103Q-FP imposes a visible growth defect to cells (**Fig. S1**). We then monitored aggregate formation in the cytoplasm of yeast cells using fluorescence microscopy, by tracking the formation of bright fluorescent foci. Cells expressing soluble proteins (i.e. with low polyQ number, such as htt19Q-FP) would not accumulate fluorescent foci, but instead would show diffuse fluorescence (58), whereas only cells overexpressing the aggregation-prone mutant of htt-FP would form clear foci (**Fig. 1D** and **Fig. S2**). We used the formation of fluorescent foci as a guide for MIP analysis.

### Wide-field fluorescence-detected MIP reveals β-sheet enrichment in protein aggregates

To characterize the secondary structure of protein aggregates, we initially fused htt protein with a green fluorescent protein (GFP). In this way, we combine the high specificity of fluorescent labeling and the structural information of mid-infrared absorption, by directly detecting MIP signals from the center of the regions of high fluorescence intensity (foci). We first harnessed GFP as a highly sensitive sensor of temperature(45) in wide-field MIP. The concept of high-throughput single-aggregate structural analysis by wide-field fluorescence-detected MIP is illustrated in **Fig. 2*A***. Wide-field fluorescence imaging of yeast cells enabled simultaneous analysis of tens of htt103Q-GFP aggregates within the mid-IR illumination area (∼50 µm in diameter) while minimizing photobleaching of GFP (45). For each single aggregate, fluorescence intensity change induced by the mid-IR absorption could be quantified. The thermal sensitivity of fluorescent proteins leads to a drop in fluorescence intensity when the temperature rises. To capture the transient change induced by the ultrashort mid-IR pulses (pulse width ∼20 ns), the fluorescence excitation is synchronized to the mid-IR pulses and modulated into pulses (width ∼200 ns). Limited by the relative slow speed of camera (∼60 ms per frame), the mid-IR pulses are chopped with 50% duty cycle to produce alternating IR on (‘hot) and IR off (‘cold’) frames. The fluorescence intensity traces of single aggregates are analyzed to calculate the MIP modulation depth (MD).

**Fig. 2.**
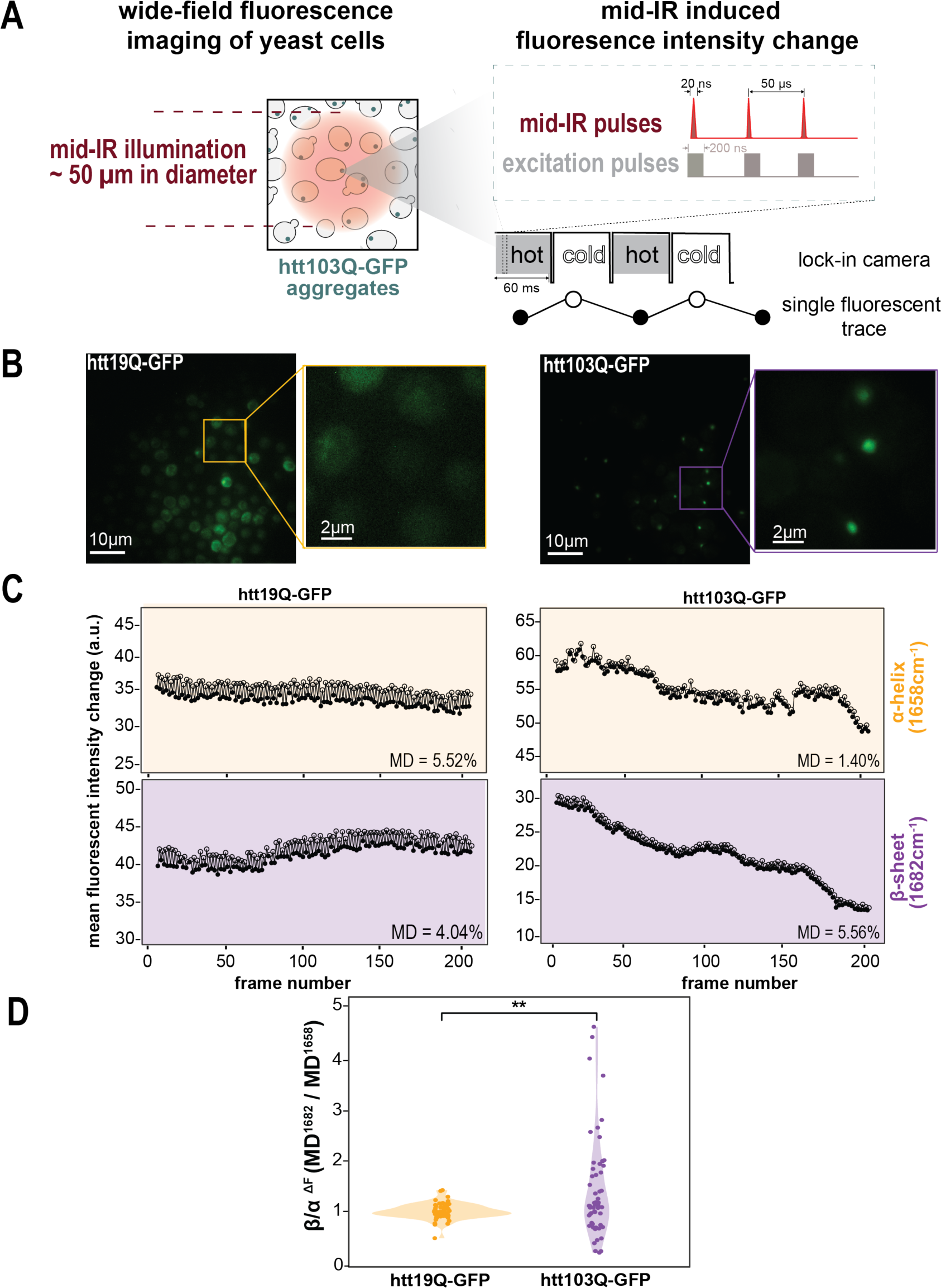
Wide-field fluorescence-detected MIP reveals β-sheet enrichment in fluorescently labeled protein aggregates with high throughput. (*A*) Schematic of high-throughput single aggregate structural analysis by wide-field fluorescence-detected MIP. Wide-field mid-IR illumination allows simultaneous analysis of multiple single aggregates. Mid-IR induces fluorescence changes, which are measured by synchronized fluorescence excitation pulses and ‘lock-in’ camera of alternating ‘hot’ and ‘cold’ frames. Fluorescence intensity traces for single aggregates were analyzed for modulation depth at each IR wavenumber. (*B*) Representative wide-field fluorescence images and detail images of GFP fluorescent trace for htt19Q-GFP (left) and htt103Q-GFP (right). To successfully visualize htt19-GFP fluorescence, the camera gain for htt19Q-GFP was 20dB higher than for htt103Q-GFP. Scale bars: 10 µm and 2 µm for detail inset. (*C*) Representative fluorescence intensity traces with mid-infrared at 1658 cm^-1^ for α-helix (top) and at 1682 cm^-1^ for β-sheet (bottom) for htt19Q-GFP cell (left) and htt103Q-GFP foci (right) at the center of the enlarged selection in images in (*C*). MD: modulation depth. (*D*) Statistical analysis of fluorescence modulation depth ratio at 1682 cm^-1^ over at 1658 cm^-1^ (β/ɑ^ΔF^) for htt19Q-GFP cell (N = 63) and htt103Q-GFP foci (N = 55). Unpaired pairwise t-test, p = 0.001 **.

To distinguish structural properties of htt aggregates, we focused on the two extreme cases of a very low polyQ mutant (htt19Q), which is mostly soluble, and a very high polyQ mutant (htt103Q), which is very aggregation prone. We imaged yeast strains expressing each mutant respectively. As expected, the wide-field fluorescence images of yeast strains expressing htt19Q-GFP showed diffuse fluorescence while htt103Q-GFP showed bright foci (**Fig. 2*B***, S2). Fluorescence videos were recorded with mid-IR excitation at 1658 cm^-1^ (representing ɑ-helix) and at 1682 cm^-1^ (representing β-sheet) sequentially. The fluorescence intensity traces from representative htt19Q-GFP cell body and htt103Q-GFP foci showed clear modulation where hot frames (solid circle) had lower intensity compared with adjacent cold frames (open circle) (**Fig. 2*C***). Therefore, the MIP modulation depth was calculated as mid-IR induced change relative to the original fluorescence intensity for single htt19Q-GFP cell and htt103Q-GFP foci. htt103Q-GFP showed higher modulation at 1682 cm^-1^ while htt19Q-GFP showed higher modulation at 1658 cm^-1^. We note that the htt19Q-GFP cell showed higher fluorescence modulation depth than htt103Q-GFP foci, presumably as a result of the effect of solvent viscosity on fluorescence intensity in addition to temperature change (59). Since the wide-field configuration is conducive to high throughput applications, we performed statistical analysis on the ratio between fluorescence modulation at 1682 cm^-1^ and at 1658 cm^-1^ (β/ɑ^ΔF^) and found a higher ratio for htt103Q-GFP compared with htt19Q-GFP (**Fig. 2*D***). This higher ratio indicates an enrichment of β-sheet structure in protein aggregates, which is consistent with *in vitro* representations of polyQ fibrils as amyloid fibrils formed by a core of interdigitates β-hairpins, or a stacked β-sheet (60, 61). Collectively, these data demonstrate that mid-IR-modulated fluorescence is able to reveal β-sheet enrichment in protein aggregates and could be used to identify amyloid aggregates.

### Fluorescence-guided MIP spectroscopy dissects secondary structure of protein aggregates

As β-sheet enrichment of protein aggregates could be detected by fluorescence-detected MIP, we combined MIP spectroscopy and fluorescence microscopy to explore the structural properties of protein aggregates in yeast cells. The schematic for fluorescence-guided MIP spectroscopy is shown in **Fig. 3*A***. Epi-fluorescence imaging were first performed to provide guidance for the intracellular location of protein aggregates. The fluorescence module can be easily switched to accommodate the need for imaging both green and red fluorophores. As the image was formed by the raster scanning of the sample stage, the sample stage could be driven by piezo voltage to target single aggregates sequentially based on the center pixel positions. Then, both mid-IR and visible probe beam were focused on to the target aggregate, the MIP spectrum was acquired by continuously tuning the mid-IR wavenumbers.

**Fig. 3.**
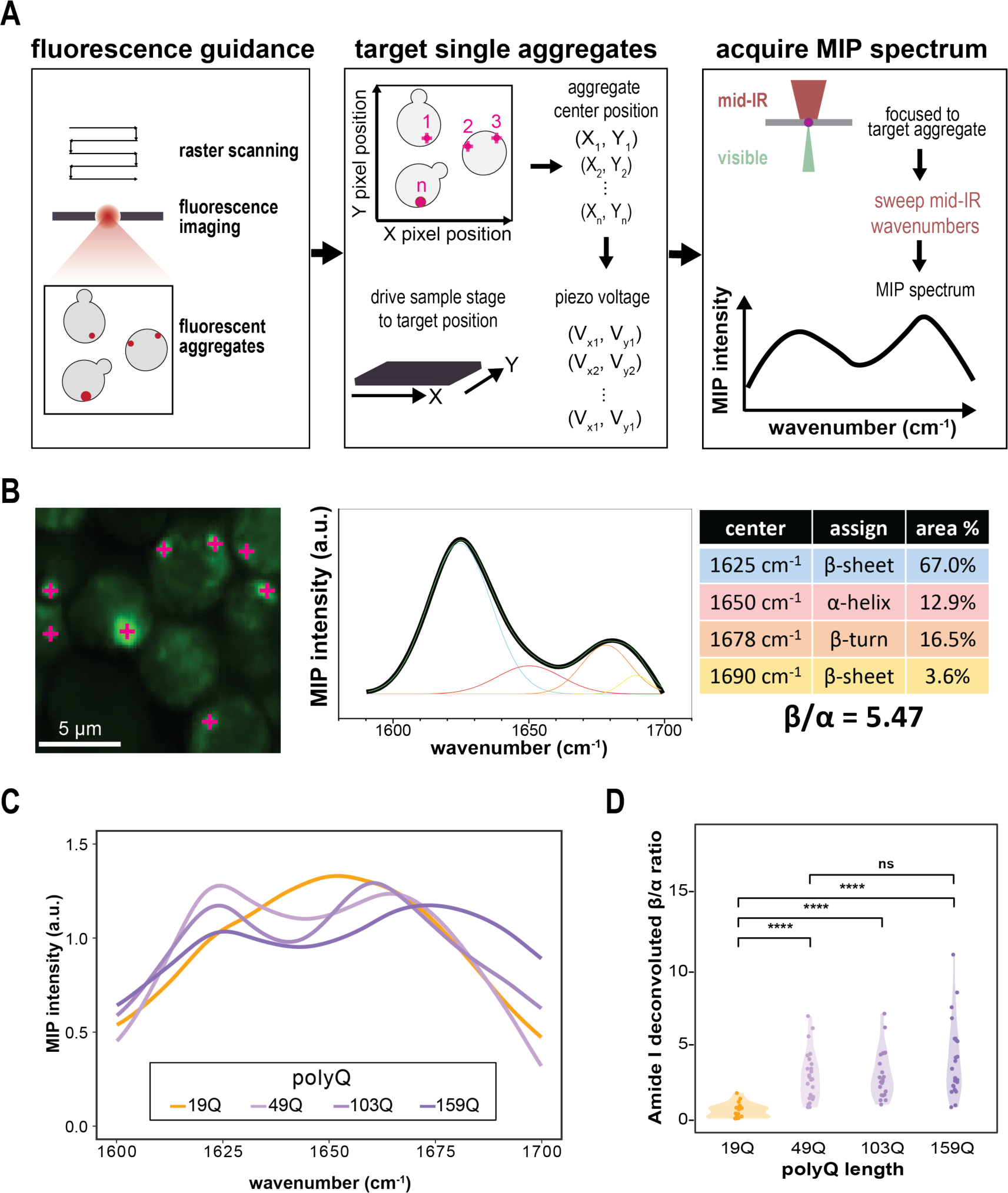
Fluorescence-guided MIP spectroscopy dissects spectral features of protein aggregates. (*A*) Schematic for fluorescence-guided MIP spectroscopy. Fluorescence imaging was first performed by raster scanning of the sample stage. To target single aggregates, the sample stage was driven by piezo voltages based on pixel positions of aggregate centers sequentially. The MIP spectra were acquired by continuously tuning the mid-IR wavenumbers. (*B*) A representative MIP spectrum of htt103Q-GFP aggregate acquired from fluorescent foci (bright green spots marked by purple crosses) and deconvoluted in amide I region into component peaks assigned to different secondary structures. β/ɑ ratio was calculated by area under the component peaks. (*C*) Representative MIP spectra in amide I region acquired from 159Q, 103Q and 49Q aggregates under Crimson fluorescent guidance, and from 19Q yeast cells. (*D*) β-sheet to ɑ-helix ratio (β/ɑ) quantified by amide I deconvolution for 19Q, 49Q, 103Q and 159Q. Unpaired pairwise t-test, N>20, p<0.0001 **** and p > 0.05 ns.

We acquired a set of MIP spectra across protein aggregates, by sequentially moving the stage to the center of fluorescent foci (bright green spots marked by purple crosses). **Fig. 3*B*** is a representative example of MIP spectroscopic analysis of protein aggregates in a field of cells expressing htt103Q-GFP. The MIP spectrum showed significant shifts towards both low and high wavenumbers, instead of a broad amide I band. MIP spectrum in the amide I region was deconvoluted into multiple component peaks, the center of which was determined by the minimums of the second derivative. Each peak could be assigned to different protein secondary structures based on their peak centers. The area under a component peak quantifies its relative contribution. We calculated the β/ɑ ratio to quantify the β-sheet enrichment as the ratio of the sum of β-sheet components peak area to that of ɑ-helix, confirming that the core of htt aggregates (defined as the center of bright foci) is rich in β-sheet structure. A detailed illustration of signal processing and spectral deconvolution analysis could be found in **Fig. S3**. We note no obvious heat accumulation due to strong infrared absorption during wavenumber sweep for MIP spectra under our experimental settings (**Fig. S4**). We also note that fixation made β-sheet structures in aggregates undetectable by MIP, possibly due to the effect of cross-link of protein by paraformaldehyde (62, 63) (**Fig. S5** and Supplementary Methods). A custom LabVIEW program was developed for interactive control (Supplementary Data).

### PolyQ length does not affect secondary structure of htt aggregates *in vivo*

By performing full MIP scanning of fluorescent foci, we aimed at exploring how the structure of htt aggregates varies across mutants. Studies *in vitro* with purified peptides demonstrated that polyQ proteins, given enough time, always converge in forming β-sheet rich (amyloid) fibrils in a repeat-length dependent fashion at a rate dictated by the polyQ length (64). This observation is in contrast with the well-established threshold effect seen in all polyQ diseases, where a specific polyQ length is considered the threshold (38 repeats for HD) above which toxicity arises and pathology develops (65). In yeast, it is well established that aggregation propensity of htt correlates with polyQ length (58), but there is yet scarce information on a correlation between polyQ length and structural properties of the aggregates in their physiological state. Hence, we were interested to study whether polyQ aggregates accumulating in cells invariably form β-sheet rich inclusions or if they vary in structural properties with polyQ length.

We expressed htt fusions with the fluorescent protein Crimson (htt*n*Q-Cr) with variable polyQ length (respectively, 19Q, 49Q, 103Q and 159Q) and we analyzed yeast cells with MIP. The representative spectra acquired from the fluorescent foci for Q length above the pathological threshold (159Q, 103Q and 49Q) showed a similar spectral change of increasing low- and high-wavenumber β-sheet, whereas 19Q showed a broad amide I band (**Fig. 3*C***). We further performed quantitative analysis of β/ɑ ratio after amide I deconvolution. Statistical analysis showed lower β/ɑ ratio for 19Q, compared with 159Q, 103Q and 49Q while no significant difference between the 3 polyQ length above the threshold (**Fig. 3*D***). Interestingly, when we extended the analysis of β/ɑ ratios to fluorescent traces of proteins with shorter polyQ, we could fit a sigmoidal curve correlating polyQ length with β/ɑ ratios (**Fig. S6**), with the case of 37Q forming foci (unlike 19Q, 26Q and 31Q) but with weaker β/ɑ ratio compared to 49Q-159Q and no difference in β/ɑ ratio above threshold. Collectively, this data is consistent with a scenario where polyQ length affects rate of aggregation, but once aggregates are formed, all acquire the same ultimate secondary structure. Slower rates of aggregation enable cells to cope with accumulating aggregates, which explains the diverse β/ɑ ratio across strains expressing short polyQs (66).

### Label-free structural study by MIP reveals secondary structure perturbation in htt aggregates caused by fusion of GFP

With MIP it is possible to scan a sample at a defined set of wavenumbers, therefore mapping chemical information onto a microscopy image. We argued that, once learned the spectral signature of a certain type of protein aggregates, it would be possible to perform label-free analysis, by imaging cells expressing htt103Q devoid of GFP. To do this, we first tested live yeast cells expressing htt103Q-GFP, by sequentially acquiring fluorescence images (in green) and MIP images at 1658 cm^-1^ (representing ɑ-helix, in red), 1628 cm^-1^ (representing low-wavenumber β-sheet, in cyan), and 1682 cm^-1^ (representing high-wavenumber β-sheet, in yellow), described in **Fig. 4*A***. We observed co-localization between GFP bright foci and strong MIP signals at both β-sheet wavenumbers (1628 and 1682 cm^-1^). This result is compatible with the large literature of *in vitro* studies that describe polyQ fibers as stacked β-sheets (67). This shows that multispectral MIP imaging at a few selected signature wavenumbers suffice to represent the spatial distribution of β-sheet and ɑ-helix structures of protein aggregates.

**Fig. 4.**
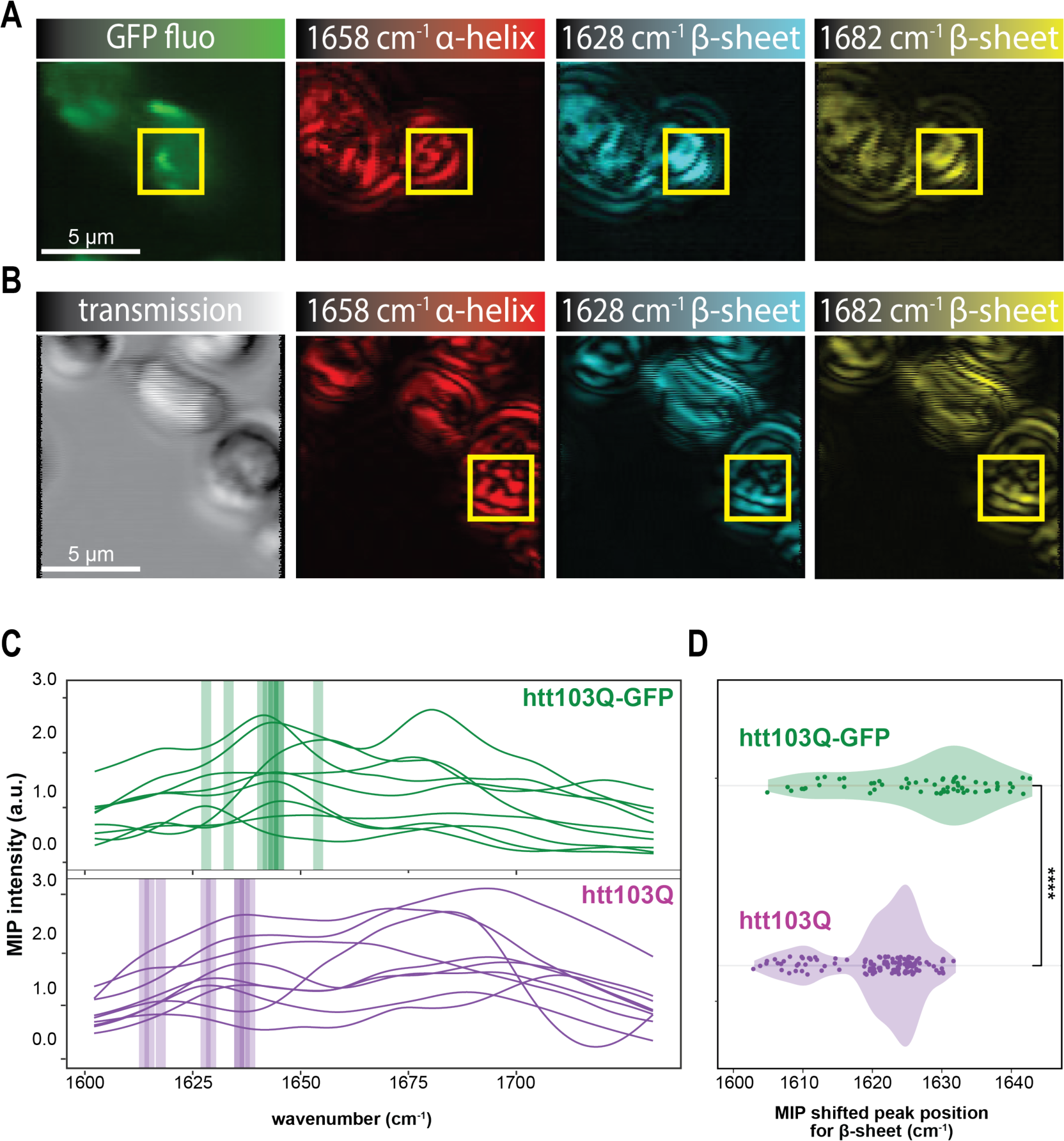
MIP detects structural perturbation caused by fusion of GFP. (*A*) Representative fluorescence images for GFP (green) and MIP images at 1658 cm^-1^ (ɑ-helix, red), 1628 cm^-1^ (low-wavenumber β-sheet, cyan), and 1682 cm^-1^ (high-wavenumber β-sheet, yellow) for live yeasts expressing htt103Q-GFP. Scale bar: 5 µm. (*B*) Representative transmission image (gray) and MIP images at 1658 cm^-1^ (ɑ-helix, red), 1628 cm^-1^ (low-wavenumber β-sheet, cyan), and 1682 cm^-1^ (high-wavenumber β-sheet, yellow) for live yeasts expressing htt103Q with no fluorescent tag. Scale bar: 5 µm. The yellow box identifies the protein aggregate in each frame of A and B. (*C*) Representative 9 MIP spectra in amide I region for htt103Q-GFP aggregates (top) with a shifted peak position for low-wavenumber β-sheet and Htt103Q aggregates (bottom) with a further red-shifted peak position. (*D*) MIP shifted peak position for low-wavenumber β-sheet for htt103Q-GFP (N = 62) and htt103Q (N = 135) aggregates. Unpaired pairwise t-test, p = 3.18ξ10^-6^, ****.

It is known that fluorescent proteins can alter the biophysical properties of a protein, by introducing steric hindrance, charge repulsion, or inducing oligomerization (68, 69). We hypothesized that the large size of GFP (double in molecular weight compared to htt) would affect the structure of htt aggregates and that a label-free modality of structural imaging would provide more faithful insights on the structure of aggregates in their physiological state.

We therefore acquired MIP images at 1658, 1628 and 1682 cm^-1^ of cells expressing htt103Q with no fluorescent protein tag (**Fig. 4*B***). The chosen wavenumbers are representative of, respectively, ɑ-helix, low-wavenumber β-sheet and high-wavenumber β-sheet. We scanned for areas characterized by high β-sheet and relatively low ɑ-helix to identify htt103Q aggregates. We observed that the MIP spectra for htt103Q aggregates identified by structural features showed a further red shifted peak position for low-wavenumber β-sheet compared with htt103Q-GFP aggregates (**Fig. 4*C***). Statistical analysis showed a further red shift (i.e. even lower wavenumber) in the shifted peak position for β-sheet in htt103Q aggregates (N = 135) compared with htt103Q-GFP aggregates (N = 62) (**Fig. 4*D***).

This contribution of the fluorescent tag can be explained in two ways: first, their size, charge and propensity to aggregate can destabilize or influence the morphology of the aggregates. Second, proteins such as GFP are distinctively enriched in β-sheets, hence potentially misleading an analysis based on secondary structure features. However, FTIR studies of purified proteins *in vitro* showed that it is possible to distinguish the contribution of native β-sheet (i.e. relative to the structure of a protein such as GFP) and that of amyloids. Indeed, native β-sheet proteins such as GFP have a shifted amide I peak position from 1630 to 1643 cm^-1^, while amyloid fibrils present the amide I peak clustering between 1611 and 1630 cm^-1^ (70). Therefore, analyzing the shift in amide I peak position provides specific information on the aggregate (71, 72). Collectively, this data suggests that tagging aggregates with large fluorescent proteins introduces confounding effects both by introducing more spectral contributions and by directly perturbing the structure of the aggregate itself.

To the best of our knowledge, this is the first time that this spectral shifted position difference was observed in protein aggregates in their physiological condition by MIP. Therefore, MIP offers a new approach to study protein aggregates without potential interference from fluorescent proteins.

### MIP spectroscopy reveals structural partitioning within protein aggregates below the resolution limit

Interestingly, by examining the details of secondary structural distribution of htt103Q-GFP aggregate (represented in **Fig 4*A***), we observed a spatial distribution of high intensities for ɑ-helix wavenumber (1658 cm^-1^) around the areas of high β-sheet intensity, where the ɑ-rich region forms a hollow-like distribution around the β-rich region (**Fig 5*A***). This suggests that htt aggregates are organized into a β-sheet enriched core, surrounded by a ɑ-helix dominant outer layer. By tapping into the additional dimension of spectral information, we could detect the aggregation core enriched in β-sheet components, despite being of a lower size than the visible probe beam (73, 74) (**Fig. S7**).

**Fig. 5.**
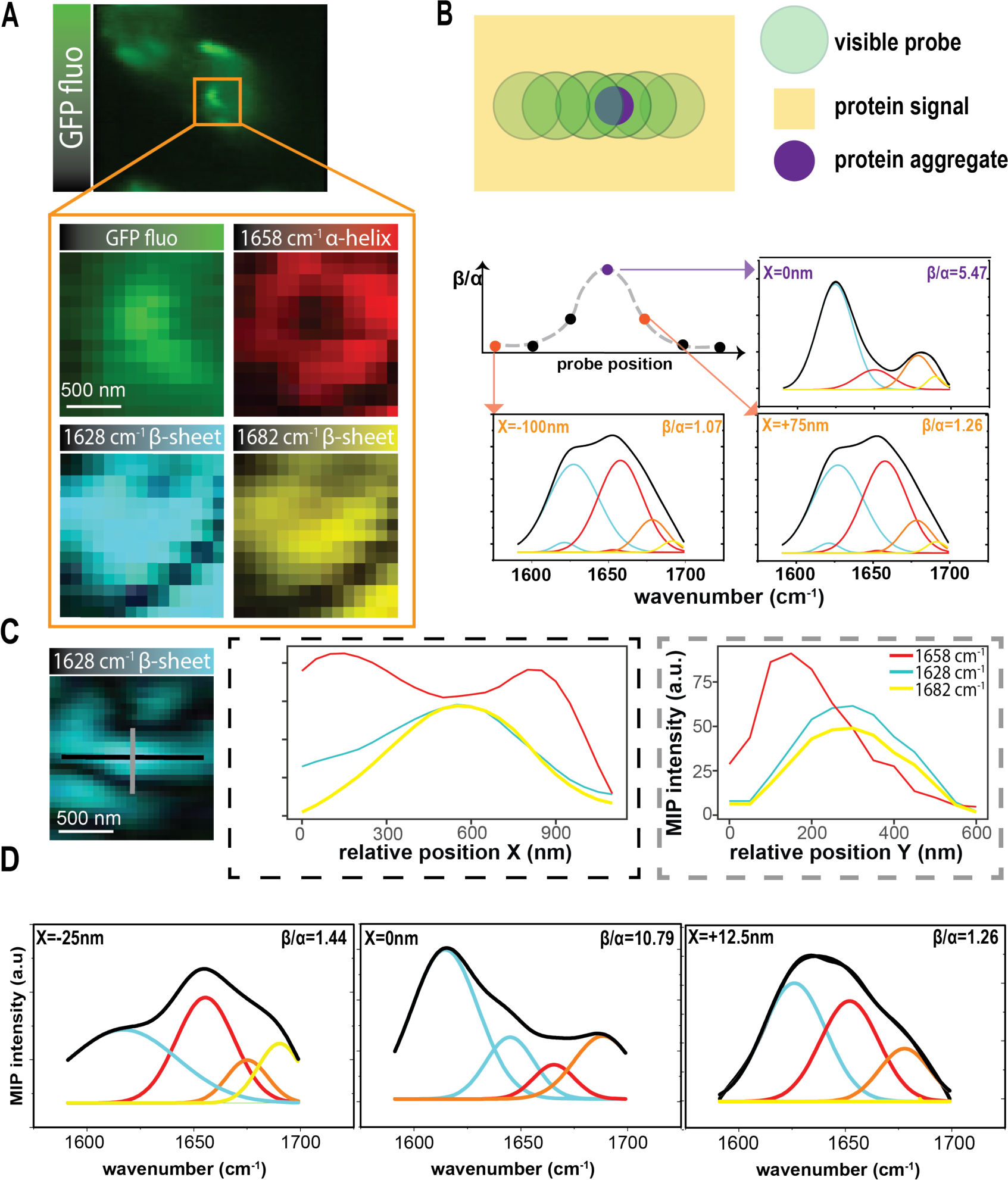
MIP resolves structural partitioning of htt aggregates at the nanometer scale. (*A*) detailed spatial distribution of htt103Q-GFP aggregate revealed a β-sheet enriched core surrounded by a ɑ-helix dominant outer layer. (*B*) MIP spectra with small spatial steps across the β-sheet enriched core revealed drastic change in β-sheet enrichment for htt103Q-GFP aggregate with MIP spectra acquired at relative positions x = -100, 0, +75 nm in amide I region (overall spectra in black, component peaks colored) and quantified β-sheet to ɑ-helix ratios (β/ɑ). For htt103Q aggregate: (*C*) X-dimension (left) and Y-dimension (right) line profile of identified aggregate at 1658 cm^-1^ (red), 1628 cm^-1^ (cyan) and 1682 cm^-1^ (yellow) and (*D*) MIP spectra acquired at relative positions x = -25, 0, +12.5 nm in amide I region (overall spectra in black, component peaks colored) and quantified β-sheet to ɑ-helix ratios (β/ɑ).

The peculiar structural partitioning of ɑ-rich and β-rich regions observed in **Fig. 5*A*** motivated us to quantitatively explore the spectral features throughout the protein aggregate. We hypothesized that this partition is intrinsic to the nature of htt aggregates. To do this, we recorded full MIP spectra sequentially along one direction across the aggregation complex with small spatial steps (**Fig. 5*B***). We noticed how spectra progressively shifted from ɑ-rich to β-rich and back to ɑ-rich along the cross-section, as showed by the variation in β/ɑ ratio along the axis (**Fig. 5*B***, **Fig. S8** and **S9**). We acquired MIP spectra from x = -100, 0, and +75 nm relative positions and quantified the β/ɑ ratio. The center showed the highest β/ɑ of 5.47 and β/ɑ decreased to 1.17 (x = +75 nm) and 1.07 (x = -100nm) away from the center. This analysis validated the observation in **Fig. 5*A***.

In order to exclude potential contribution of GFP to this partitioning, we performed small step full MIP spectra also on htt103Q aggregates (devoid of GFP fusion). We plotted the X-dimension and Y-dimension line profile for this identified aggregate at 1658 cm^-1^, 1628 cm^-1^ and 1682 cm^-1^ and observed a center with high β-sheet and corresponding dip in ɑ-helix (**Fig. 5*C***). Once we acquired MIP spectra with small spatial steps, we observed an even higher β-sheet to ɑ-helix ratio (β/ɑ) of 10.79 at the aggregation center as well as a more drastic drop in β/ɑ ratios 25 and 12.5 nm away from the center (**Fig. 5*D***), suggesting that this spatial partitioning is not caused by GFP.

Collectively, we revealed the spatial distribution of structurally distinct components within the protein aggregates through direct visualization by multispectral MIP imaging at signature wavenumbers. Additionally, by moving along the aggregate body with nanometer sized spatial steps, we revealed different degrees of relative contribution of the β-sheet core to the acquired MIP spectra, causing drastic spectral change within small distances below the conventional optic diffraction limit. This demonstrates how performing quantitative analysis of full MIP spectra with small spatial steps can dissect the protein aggregates into distinct structural regions.

### MIP micro-spectroscopy reveals the influence of the Rnq1 prion on the structural properties of htt aggregates

In our micro-spectroscopy experiments, we observed spatial partitioning of different secondary structures within htt aggregates (**Fig. 5**). We hypothesized that this structural heterogeneity is a consequence of the heterogeneous composition of the aggregates. The prion protein Rnq1 is known to aggregate in a prion form known as [*RNQ^+^*]. This protein has unknown functions when soluble, but, in its prion form, it is known to facilitate formation of other prions, such as the one of the protein Sup35 (i.e. the prion state [*PSI^+^*]) (75). Indeed, it is also well-known that the [*RNQ*^+^] state can facilitate htt aggregation in *S. cerevisiae* and induces toxicity (76). In fact, there is evidence of co-localization of htt aggregates and Rnq1 protein by fluorescent microscopy (77). Moreover, recent work has described the formation of dynamic condensates by htt in yeast (78, 79), and it is known that condensates formed by aggregation-prone proteins follow a maturation process (vitrification) where they progressively transition from dynamic liquid-like droplets into solid-like fibrillary structures from within the droplet (80–82). We therefore hypothesize that the structural partitioning described in **Fig. 5** is caused by the heterogeneous condensation of Rnq1 protein, which is an intermediate step towards formation of fully fibrillary aggregates.

To investigate this, we used a recently developed system to induce polyQ aggregation in [*rnq^-^*] cells that would otherwise not form consistent polyQ aggregates (47) (**Fig. 6*A***). Here, htt was fused to Cry2 (and the mCherry fluorophore), a protein derived from *Arabidopsis thaliana* that is widely used in optogenetics experiments and known to induce oligomerization in response to blue light stimulation (83). We generated yeast strains containing inducible versions of htt43Q-Cry2-mCherry in either a [*RNQ*^+^] or [*rnq*^-^] background. To incorporate blue light stimulation into our microscope, we used the powerful illumination lamp in the MIP microscope and added a band pass filter (480/10 nm) before focusing the blue light on to the sample by the IR objective.

**Fig. 6.**
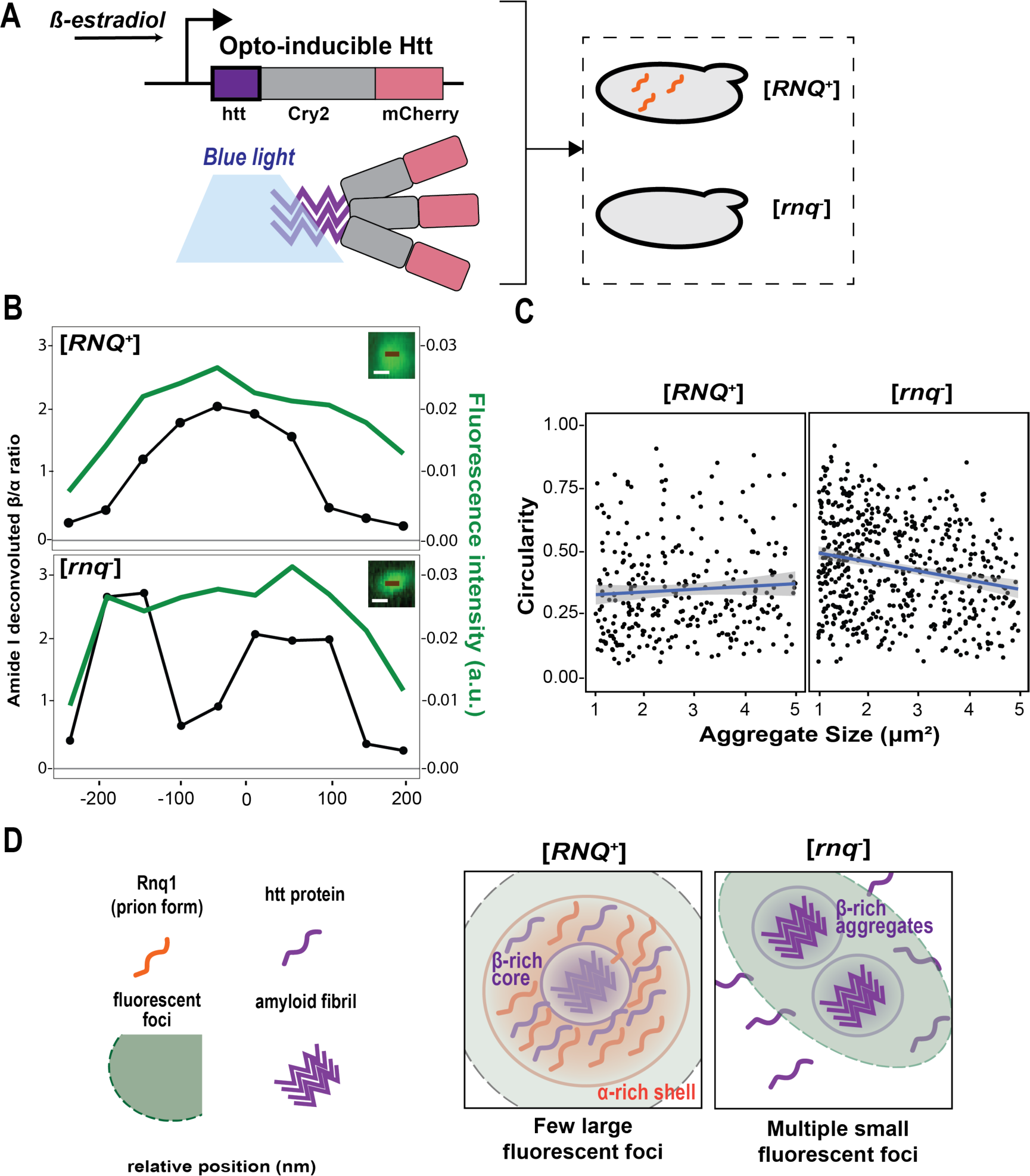
MIP detects perturbations caused by the prion Rnq1 on the structural properties of htt aggregates. (*A*) Experimental approach to study the role of prion Rnq1 in htt aggregation. In both [*RNQ^+^*] and [*rnq-*] cells, β-estradiol induces expression of recombinant htt43Q-Cry2-mCherry and blue light stimulates nucleation of the aggregate (*B*) β/ɑ ratio plot along the line of protein aggregates overlaid with fluorescence intensity for [*RNQ^+^*] strain (top) and [*rnq*^-^] strain (bottom). Inset: mCherry fluorescence images in green, scale bar 500 nm. (*C*) Circularity of fluorescent foci against the aggregate size in [*RNQ*^+^] (left) and [*rnq^-^*] strains. Each dot represents single fluorescent foci. Blue line shows linear fitting. (*D*) Schematic representation of htt aggregates in [*RNQ^+^*] and [*rnq^-^*] cells: the prion form of Rnq1 protein (orange) nucleates htt aggregation (purple), by forming α-rich aggregates, from which amyloid (β-rich) fibrils maturate, leading to one large fluorescent foci. In [*rnq^-^*] cells, aggregation of htt (when possible) follows a separate aggregation kinetics than in [*RNQ^+^*] cells, bypassing the α-rich nucleation step, leading to multiple smaller and β-rich fluorescent foci.

We observed formation of new fluorescent foci after blue light stimulation in estradiol-induced Cry2-mCherry cells in both [*rnq*^-^] and [*RNQ*^+^] backgrounds (**Fig. S10**). We then expressed a htt43Q-Cry2-mCherry fusion and observed that htt43Q-Cry2-mCherry successfully formed aggregates in both backgrounds, with the average number of fluorescent foci increasing after blue light stimulation for 5 min (**Fig. S11**). No difference in the β/ɑ ratio from MIP spectra was observed between htt43Q-Cry2-mCherry and htt49Q-Crimson in [*RNQ^+^*] strains (**Fig. S12**), suggesting that Cry2 does not perturb the type of aggregates formed, but only induces their nucleation.

We then compared MIP spectra and β/ɑ ratios across the htt43Q-Cry2-mCherry aggregates in [*RNQ^+^*] and in [*rnq^-^*] strains. MIP analysis of fluorescent foci in [*RNQ^+^*] showed symmetric single peaks in the β/ɑ plots of their cross-section (with a 50 nm step), as expected, whereas large bright fluorescent foci found in [*rnq^-^*] presented fluctuations with multiple peaks in the β/ɑ plots of their cross-section (**Fig. 6*B***, **Fig. S13**). This suggested the presence of multiple smaller aggregates close in space, but below the diffraction limit.

We hypothesized that, in this scenario, large bright foci in [*RNQ^+^*] would have a round shape, whereas large bright foci in [*rnq^-^*] would have lower circularity due to the combined fluorescence of two sources close in space. Hence, circularity of aggregates should scale with size in the [*rnq^-^*] background, but not scale in the [*RNQ^+^*] background. To validate this, we plotted circularity against particle size of aggregates within cells. Indeed, particle size had an inverse correlation with circularity value in aggregates found in [*rnq^-^*], whereas those found in [*RNQ^+^*] had no variation of circularity across sizes (**Fig. 6*C***). Finally, we measured cell growth curves in presence of blue light for yeast cells expressing htt43Q-Cry2-mCherry in both backgrounds (**Fig. S13**). Interestingly, growth in [*RNQ^+^*] was slower compared with [*rnq^-^*] strains (**Fig. S14**), confirming that the presence of the prion form of Rnq1 protein is associated with toxicity in this cellular model of Huntington’s Disease (47).

Collectively, we demonstrate that MIP spectroscopy is capable of detecting ultrastructural perturbations caused by the prion form of Rnq1 on htt aggregates. Our results suggest that the Rnq1 prion protein is not only associated with the nucleation of the aggregates, but influences the physical and structural properties of the aggregate. We propose that, when nucleated by Rnq1 amyloid, htt aggregates are initially dynamic, mostly composed of proteins in α-helical conformation and with properties close to those of liquid droplets (79), but then quickly progress into more rigid and fibrillary structures. The core-shell distribution is therefore reminiscent of this transition (81). Conversely, in absence of other prions, htt forms smaller, exclusively β-sheet and potentially non-miscible aggregates, suggesting the nucleation step is followed directly by fibril formation (**Fig. 6*D***).

## Discussion

Protein aggregation and, more specifically, amyloid formation are key hallmarks of multiple neurodegenerative diseases, such as Alzheimer’s and Parkinson’s Disease (84, 85), and the direct cause of neurodegeneration in Huntington’s Disease (HD) and other disorders of the family of the polyglutamine diseases (86–88). Disease-relevant proteins, such as the huntingtin protein in HD, accumulate intracellularly and are known to form large, ordered structures composed of stacked polypeptides in β-sheet conformation to form a fibril. Amyloid structures have been thoroughly characterized *in vitro* via multiple techniques, including NMR (89) and cryo-EM (18, 90). Using such methods, purified huntingtin has been described *in vitro* to form fibrils by stacking interdigitated β-hairpins along the fibril axis (60, 91).

Despite the wealth of structural information on aggregates *in vitro*, it is challenging to translate these structural insights to the physiological and cellular state of protein aggregates. The only accounts of aggregate structure within cells have been obtained using cryo-EM and super-resolution microscopy. For instance, with cryoEM has been possible to obtain high resolution structure of MAPT fibrils in Alzheimer’s Disease patients (92). Likewise, super resolution microscopy techniques such as PALM contributed in describing the morphology of intracellular aggregates (93). These approaches, however very powerful, are limited due to the technical complexity of the instruments used or their labor intensiveness that prevent the analysis of multiple cells at a time.

In this work, we used MIP to reveal the spatial distribution of structurally distinct components within protein aggregates of a cellular model of the neurodegenerative disease HD. We generated *S. cerevisiae* strains expressing polyQ-expanded variants of the HD-associated huntingtin (htt) protein. *In vitro*, polyQ proteins form aggregates at a rate dependent on their polyQ length and their final aggregated form is that of an amyloid fibrils (91, 94). However, this length dependence does not directly translate in the context of a cell. Indeed, expansion mutations in the polyQ region of htt exon 1 introduce a toxic gain of function and produce visible aggregates in the cell, only when the repeat length exceeds the threshold of 38 repeats (95); however, information on the structure of these aggregates is fragmented (96). The discrepancy between *in vitro* data and phenotypic behaviors of htt protein, and polyQ proteins more generally, motivate the development of methods to study the structure of these aggregates in their physiological state.

Here, we show that yeast cells over-expressing htt proteins with high polyQ number (49Q-159Q) accumulate aggregates that are characterized by β-sheet conformation, confirming their fibrillar nature in live cells (**Fig. 3**). To do this, we demonstrated, for the first time, that MIP can be applied to study structures of protein aggregates in live cells. Previous reports have described MIP imaging of cells during cell division, where the protein content was monitored by a single wavenumber of 1650 cm^-1^ (35). We used multiple wavenumbers in the amide I region to represent different secondary structures of protein for direct visualization of structural maps of protein aggregates.

The inherently high compatibility between MIP and fluorescence imaging enabled us to combine the rich chemical information of MIP with the high specificity of widely available fluorescent tools (97). The direct detection of MIP signals from the fluorescence intensity enabled by the wide-field configuration and synchronized fluorescence excitation provides a highly specific method to study the labeled targets with no background. Therefore, we first acquired fluorescence images and then used the fluorescent trace to identify targets of interest for MIP scanning.

We were aware that introducing a fluorescent protein tag as a fusion to our proteins of interest would likely affect the aggregation process, due to the relatively large size of the fluorescent protein tag compared to the aggregation-prone target protein, the intrinsic propensity of fluorescent proteins to oligomerize (68, 69), and the inherent structural signature of the fluorescent protein (70). Indeed, we observed further red-shifted amide I peak positions for htt103Q relative to htt103Q-GFP, which suggests that GFP indeed causes a structural perturbation. Therefore, MIP provides an alternative approach for protein aggregation studies based on spectral features and could potentially be used for diagnostics and label-free structural studies.

Once MIP was established as a *bona fide* method for analyzing secondary structure of protein aggregates in yeast cells, we interrogated mutants of htt with a range of polyQ lengths. As expected, only mutants at threshold (37Q) or above threshold (49Q-159Q) showed aggregates. Interestingly, β-sheet enrichment was equivalent in all aggregates from mutants above threshold, whereas htt-37Q showed an interesting intermediate state, with a lower β-sheet content within the aggregates, but still higher than the background values obtained by scanning cells expressing htt-19Q, htt-26Q and htt-31Q (**Fig. S6**). This is consistent with a scenario where amyloid fibrils are a stable end point of a maturation process that occurs at different rates, where this rate is dictated by the polyQ length (98, 99).

Finally, it has been proposed that htt undergoes a phase transition where its first step of condensation is the formation of liquid droplets which then evolve into their fibrillar state (100). Based on our MIP analysis, several htt aggregates present a distinct enrichment in ɑ-helical content at the edges of the β-sheet rich region within the fluorescent foci (**Fig. 5**), suggesting a combination of secondary structures arising within the aggregate. This is in agreement with a representation of htt aggregates as a fibrillar core, rich in β-sheet, surrounded by a more dynamic and liquid shell, where proteins tend to acquire a more ɑ-helical structure. Given the polymorphic nature of polyQs (101, 102) and the relatively high helical content of htt exon 1 (103, 104), it is conceivable that htt fibrils maturate from within the condensate, giving rise to the partition of structures we observed (81, 105). In this process, the amyloid form of the yeast Rnq1 protein ([*RNQ*^+^] prion state) has been proposed to facilitate nucleation of htt aggregates (55), and htt aggregates formed in the absence of this Rnq1-nucleating effect are not toxic and tend to be less liquid-like (47).

In support of this, we showed that the structural partitioning within htt aggregates disappears when htt is optogenetically-induced to aggregate in the absence of the Rnq1 prion ([*rnq^-^*] strains) (**Fig. 6**). In this condition, the aggregates observed had multiple β-rich regions within single large foci and lacked of circularity. Since the lack of circularity is an effect of objects close in space, but of sizes below the diffraction limit, we concluded that aggregates induced in [*rnq^-^*] strains tend to be smaller in size and unable to mix (**Fig.6*D***).

Altogether, we showcased how MIP can be used for detailed structural dissection of protein aggregates in live cells, elucidating the effects of polyQ length and prion composition on the secondary structure of htt aggregates.

This work unlocks the potential to expand to other neurodegenerative disease models, including primary neurons and brain tissues and to report on changes of the structural properties of neurotoxic aggregates throughout phenomena such as nucleation or liquid-liquid phase separation. Adapting this approach to proteins that are less well-characterized than htt would require complementation with biochemical, genetic and biophysical experiments, guided by MIP results.

In the future, through continued advancement in MIP instrumentation (106), it should be feasible to add a temporal dimension to MIP applications for structural kinetics studies in cells. Lastly, MIP applications could also be extended to visualize other biomolecules, such as lipids and RNAs (107, 108), using their respective characteristic infrared absorption peaks, and enabling future applications where imaging of this rich chemical complexity could be acquired in real time.

## Materials and Methods

### Plasmid assembly and cloning protocol

All plasmids generated in this study were assembled using Golden Gate, according to the assembly protocol developed for the Yeast Toolkit (56). Part 0 plasmids for htt exon 1 were generated by PCR amplification of plasmid p416 103Q GAL (58), kind donation of the Lindquist lab. Coding region for mCherry-Cry2olig was custom synthesized by Genscript, codon optimized for yeast expression. All cloning was performed in TOP10 *E. coli* cells and transformed cells were plated for selection on Lysogeny Broth (LB) agar media plates containing the appropriate antibiotic for each step of selection (chloramphenicol, carbenicillin, kanamycin). PolyQ length libraries were obtained by picking multiple positive colonies, purifying plasmid DNA and sequencing inserts to identify each repeat length. The constructs for recombinant htt43Q-Cry2-mCherry was custom synthesized by Genscript and subsequently cloned with Golden Gate as described. All plasmids used for conditional expression of the transgene were designed for chromosomal integration in the URA locus (i.e., containing 5′ and 3′ genomic homology regions and lacking a yeast origin of replication).

### Strains and growth media

The *S. cerevisiae* strains used in this work are YJW584 ([*RNQ^+^*])(109) for **Fig. 1-5** and YJW509 ([*rnq^-^*])(109) in **Fig. 6**. In both cases, background strain is W303. Strain YJW509 was obtained by curing strain YJW508 (not used in this study) with guanidine(54). Both strains were obtained by kind donation of Jonathan Weissman. Unless otherwise stated, cells were grown in synthetic medium with URA dropout (SD-URA) media at 30°C in a shaking incubator.

### Yeast transformation

Yeast colonies were grown to saturation overnight in YPD and then diluted to OD_600_ = 0.1 in 10 mL of fresh medium. Cultures were grown for approximately 6 h to OD_600_ = 0.6–0.8. Meanwhile, the transformation mixture was prepared by mixing 34 μl of DNA digestion mixture, 36 μl of 1 M lithium acetate (Sigma), 50 μl of salmon sperm DNA (Invitrogen), and 240 μl of 50% w/v PEG 3350 (Fisher). For enabling integration of the plasmid in the appropriate locus, plasmids were linearized with NotI for 1 h prior to transformation to stimulate homologous recombination. A 34 μl aliquot of this reaction mixture, without DNA cleanup, was used in the transformations. Lower amounts of DNA (up to 1:10 dilution) were used for more toxic constructs, such as those with high polyQ number. Cultures were pelleted and washed twice with sterile DI water. Washed cultures were resuspended in 1 ml of sterile DI water and then divided into 100 μl aliquots and spun down. The transformation mixture was used to resuspend pelleted cells and then incubated at 42 °C for 35 min. When selecting for prototrophy, the transformation mixture was spun down, resuspended in synthetic medium with URA dropout, and plated directly on solid agar plates (SD-URA media).

### β-estradiol induction system

The β-estradiol induction system was described previously (56). Briefly, plasmids are designed to host one single insertion cassette in the URA locus of *S. cerevisiae*. In this cassette, two genes are expressed: the first gene is a fusion of a synthetic zinc-finger transcription factor (synTF) (110) with the C-terminal ligand binding domain of Estrogen Receptor (ER-LBD) and a transactivation domain. The second gene is the gene of interest (in this work, those coding for each version of the htt protein described) under control of a synthetic promoter, where a sequence orthogonal to the yeast genome is adjacent to a minimal promoter. The synthetic sequence is specifically recognized by the synTF, such that binding of β-estradiol to ER-LBD displaces a chaperon and enables the fusion protein to translocate into the nucleus, activating the expression of a downstream gene.

Induction experiments were performed in synthetic media with 20% w/v glucose (Sigma), 0.67% w/v Yeast Nitrogen Base without Amino Acids (Sunrise Science Products), 0.2% w/v Dropout Mix without Yeast Nitrogen Base Minus Appropriate Amino Acids (Sunrise Science Products). Hormone inductions were performed with β-estradiol (Sigma).

### Yeast sample preparation for MIP imaging

The engineered yeast cells were cultured in SD-URA medium for around 62 hours after β-estradiol induction. After collection, the yeast cells were centrifuged (up to 11,000 rpm for 10 mins). About 1-2 µl of concentrated yeast cells were sandwiched between the special IR substrate and the coverglass. For wide-field fluorescence-detected MIP, silicon substrate (silicon 2018, University Wafers) was used on bottom for IR transmission and visible reflection, and coverglass on top. For point-scanning MIP, the IR-transparent calcium fluoride (CaF_2_) (0.5 mm thick) was used on top and coverglass on bottom.

### Wide-field fluorescence-detected MIP microscopy

For the wide-field fluorescence-detected microscope, the mid-infrared laser (Firefly-LW, M Squared Lasers) with a repetition rate of 20 kHz and pulse duration of about 10 ns was used as the pump. The IR power was about 43.6 mW at 1656 cm^-1^ and 42.8 mW at 1682 cm^-1^ before the microscope. The IR pulses were modulated by the optical chopper (MC2000B, Thorlabs, MC1F2 blade) with a 50% duty cycle. A 488 nm laser (06-MLD, 488nm, maximum output 200 mW) was used as the excitation source, which was digitally modulated to produce pulses of 200 ns with the laser power output set at 30 mW. For fluorescence imaging of GFP, a 500 nm short pass filter was used for excitation, a band pass filter of 525/30 nm was used for emission, along with a dichroic beam splitter (405/488/532/635 nm lasers Brightline@, quad-edge, Semrock). A 60x water objective with NA of 1.2 was used (UPlanApo, Olympus). A camera (grasshopper3 GS3-U3-51S5M, FLIR) was used to acquire the fluorescence movies at a frame rate of 16 Hz and 60 ms exposure time for each frame. A gain of 20dB was used for htt103Q and 40dB for htt19Q yeast cells. The IR pulses, fluorescence excitation pulses and the camera were all synchronized using a delay pulse generator (9524, Quantum Composers).

To avoid artifacts introduced by the movement of yeast cells, single particle tracking was performed using ImageJ plugin TrackMate (111). The mean fluorescent intensity traces were measured on tracked single particles for foci of htt103Q and cell body of htt19Q cells. The single hot-cold pair modulation depth was calculated as mid-IR induced change relative to the original fluorescence intensity, and the overall MIP modulation depth was the average with outliers removed. The MIP modulation depth was normalized by respective IR powers.

### Point-scanning counter-propagating MIP microscope

A pulsed mid-infrared pump beam (1 MHz repetition rate, 100 ns pulse duration), provided by a quantum cascade laser (QCL) (Daylight Solutions, MIRcat-2400), was focused on the sample with a reflective objective lens (40x, 0.5NA, LMM440x-P01, Thorlabs) from the top. A continuous-wave 532 nm laser (about 29mW before objective) was focused by a water immersion objective (60x, 1.2NA, UPlanSApo, Olympus) from the bottom. The forward probe photons were separated using a visible/IR dichroic mirror (GEBBAR-3-5-25, Andover Corporation), and collected onto the photodiode (PD, DET100A2, Thorlabs). The signals were electronically amplified through a 50 Ω amplifier (T4119, Thorlabs), high pass filter (0.23-1000 MHz, ZFHP-0R23-S+, Mini-Circuits), two low noise amplifiers (gain = 46 dB, 1 k – 100 MHz, SA-230F5, NF corporation) and a low pass filter (BLP-1.9+, DC-1.9 MHz, 50 Ω, Mini Circuits). The amplified signal was demodulated by a lock-in amplifier (HF2LI, Zurich Instruments). The MIP images were acquired by scanning the sample stage (scanning piezostage, Nano-Bio 2200, Mad City Laboratories) with a pixel dwell time of 100 µs and a step size of 100 nm, except for the detail images in **Fig. 5*A*** (500 µs, 50 nm).

### Fluorescence imaging on the same MIP microscope

We integrated epi-fluorescence imaging into the MIP microscope. For red fluorophores like Crimson, the probe laser 532 nm (Samba 532 nm, Cobolt) also doubles as the excitation source. A dichroic mirror with a cutoff of 550 nm (DMSP550R, Thorlabs), and emission filters of 550 nm long pass (FEL0550, Thorlabs) and 609/34 nm band pass (ET609/34m, Chroma) were used. For green fluorophores like GFP, an additional 488 nm laser was co-aligned with the 532 nm laser. We changed to a different dichroic mirror (DMSP505R, Thorlabs) and emission filter (bandpass filter 525/30 nm, Semrock) by a magnetic switchable tube. The epi-fluorescence photons were collected to a photomultiplier tube (PMT, H10721-110, Hamamatsu).

### MIP spectrum acquisition, processing, and structural analysis

To acquire MIP spectra, the sample piezo stage was moved to the target points of interest based on the voltage level corresponding to the pixel position in MIP images. The mid-IR laser will continuously sweep the wavenumbers from 1400-1780 cm^-1^ at a speed of 20 cm^-1^/s. **Fig. S3** illustrates the processing and structural analysis of MIP spectrum. The sharp dips from water vapor absorption were removed by asymmetric least square fitting assuming negative peaks. The raw MIP spectra were normalized by IR power measured by mercury cadmium telluride (MCT) detector (PVM-10.6, VIGO System). The MIP spectra in the amide I region could be considered as a linear sum of the fundamental secondary structural elements (38). The MIP spectra were averaged to 0.1 cm^-1^ and the center of component peaks were determined by the minimums of the second derivative. The component peaks were assigned to different secondary structures and quantified by the area under the peaks. The β-sheet to ɑ-helix ratios were calculated as the sum of all β-sheet peaks to that of ɑ-helix. The analysis of MIP amide I spectra were performed using OriginLab and custom-written R scripts.

### Statistical analysis

All statistical analysis was performed with unpaired pairwise t-test using the rstatix package in custom-written R scripts.

### Funding

This work was supported by the US National Institutes of Health (NIH) (grant nos. R35GM136223 and R01AI141439 to JXC), the BrightFocus Foundation (AN2020002 to ASK), Department of Defense Vannevar Bush Faculty Fellowship (no. N00014-20-1-2825 to ASK), and Schmidt Science Polymath Award (no. G-22-63292 to ASK).

## Notes

### Competing Interest Statement

The authors have declared no competing interest.

