## Supplementary Materials for "Nanoscale Structural Mapping of Protein Aggregates in Live Cells Modeling Huntington’s Disease"

### Supplementary Materials for
### Nanoscale Structural Mapping of Protein Aggregates in a Live Cells Modeling Huntington's Disease

Zhongyue Guo[1,2+], Giulio Chiesa[1,3+*], Jiaze Yin[2,4], Adam Sanford[1,3], Stefan Meier[1], Ahmad S. Khalil[1,3,5], Ji-Xin Cheng[1,2,4*]

[1]Department of Biomedical Engineering, Boston University, Boston, MA 02215, USA.
[2]Photonics Center, Boston University, Boston, MA 02215, USA.
[3]Biological Design Center, Boston University, Boston, MA 02215, USA.
[4]Department of Electrical and Computer Engineering, Boston University, Boston, MA 02215, USA.
[5]Wyss Institute for Biologically Inspired Engineering, Harvard University, Boston, MA 02215, USA.
[+]These authors contributed equally.

**Supplementary Figures**
**Supplementary Methods**
**Reference**

**Supplementary Figures**

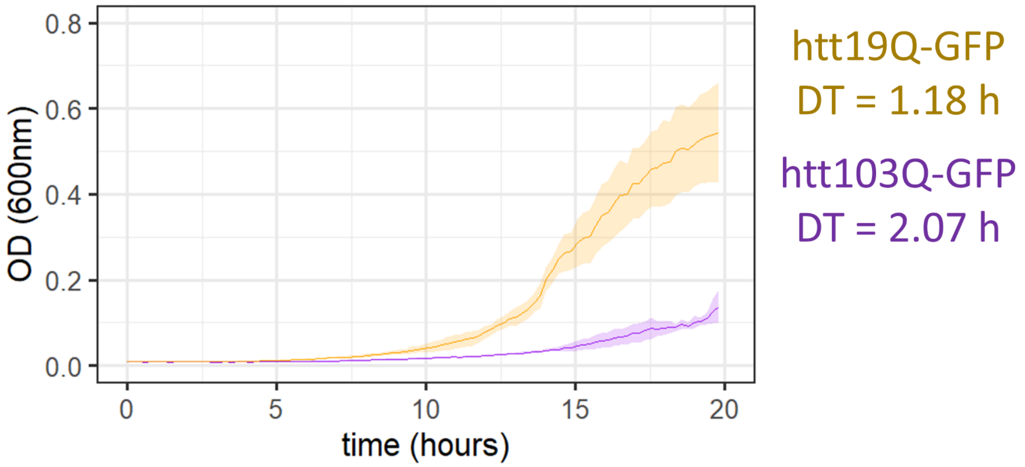


**Fig. S1.** Growth curve by absorption of 600 nm light (OD 600nm) for engineered strains with 100 nM β-estradiol induction. htt103Q-GFP (in purple) showed a slower growth compared with htt19Q-GFP (in orange). DT: doubling time. The means were represented as solid line and the standard deviations for 6 replicate wells were represented as shaded area.

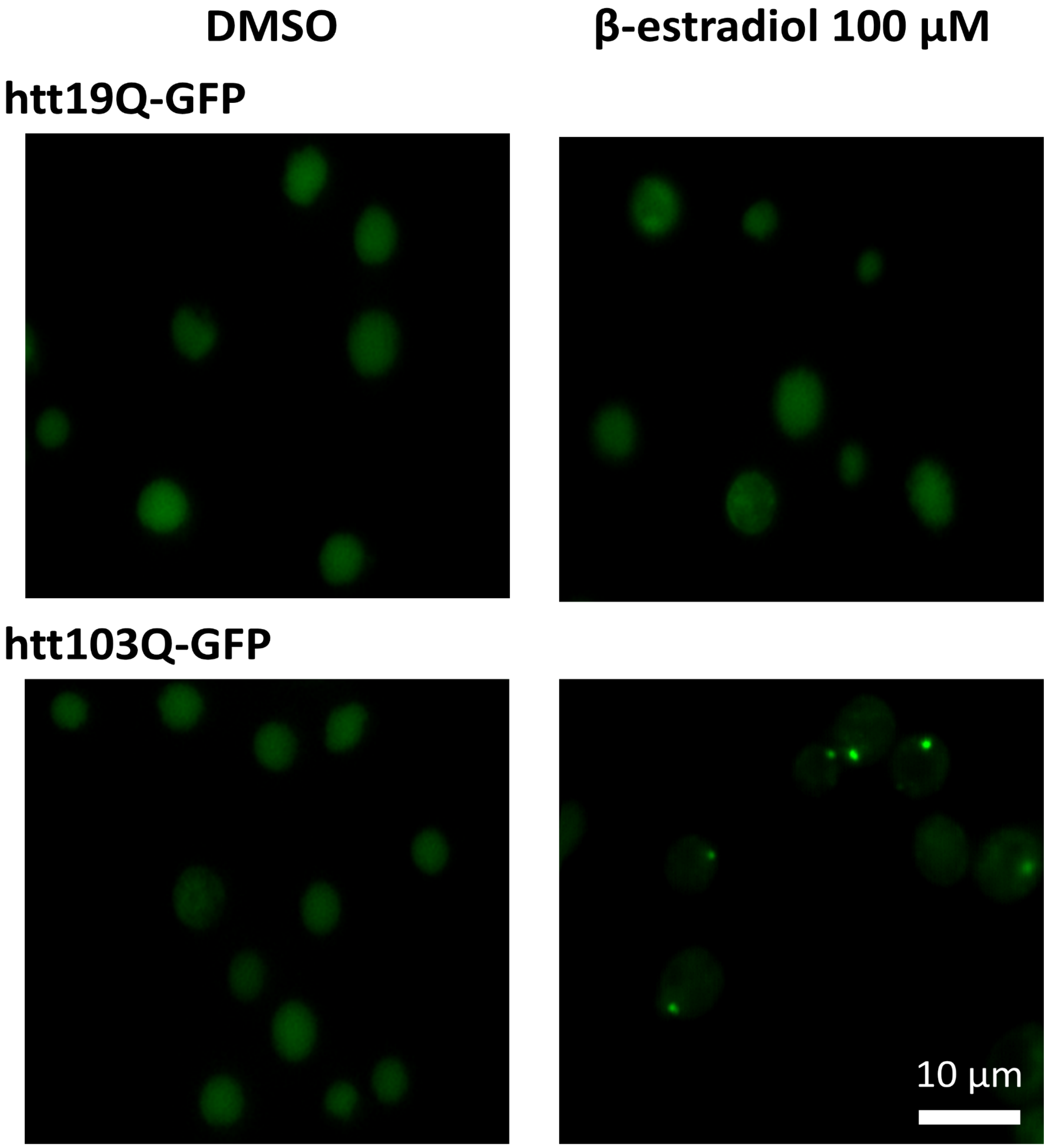


**Fig. S2**. β-estradiol induction system for modeling Huntington's disease in yeasts confirmed by fluorescence imaging. 100 μM of β-estradiol was used to induce the expression while DMSO served as control. Only induced htt103Q-GFP show clear foci while other conditions show diffuse fluorescence. Scale bar: 10 μm.

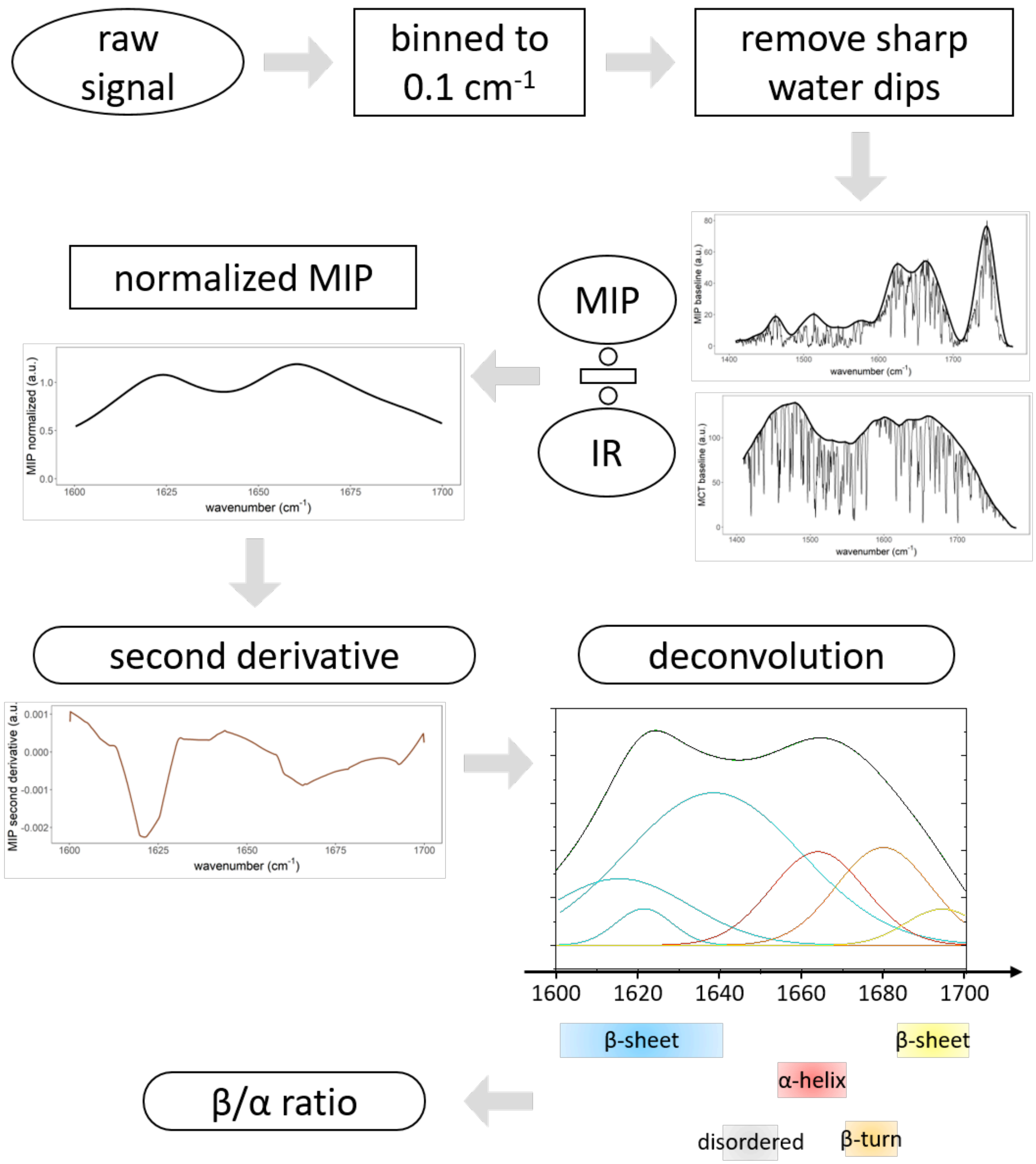


**Fig. S3**. MIP signal processing and spectral analysis. The acquired MIP raw signal was first binned to 0.1 $cm^{-1}$ spectral step. To remove the sharp water dips from absorption of mid-IR by water vapor in the environment, the baseline for each MIP spectrum and MCT detector measurement for IR power was found by asymmetric least square fitting assuming negative peaks. The MIP signal was normalized by IR power. The second derivative of normalized MIP signal was plotted in color brown. The minimum of second derivative suggested the center of hidden component peaks. The overall MIP spectrum in amide I region was fitted into component peaks and assigned to different secondary structures. Finally, the β/α ratio was calculated as the sum of all β-sheet peaks to that of α-helix.

A

B

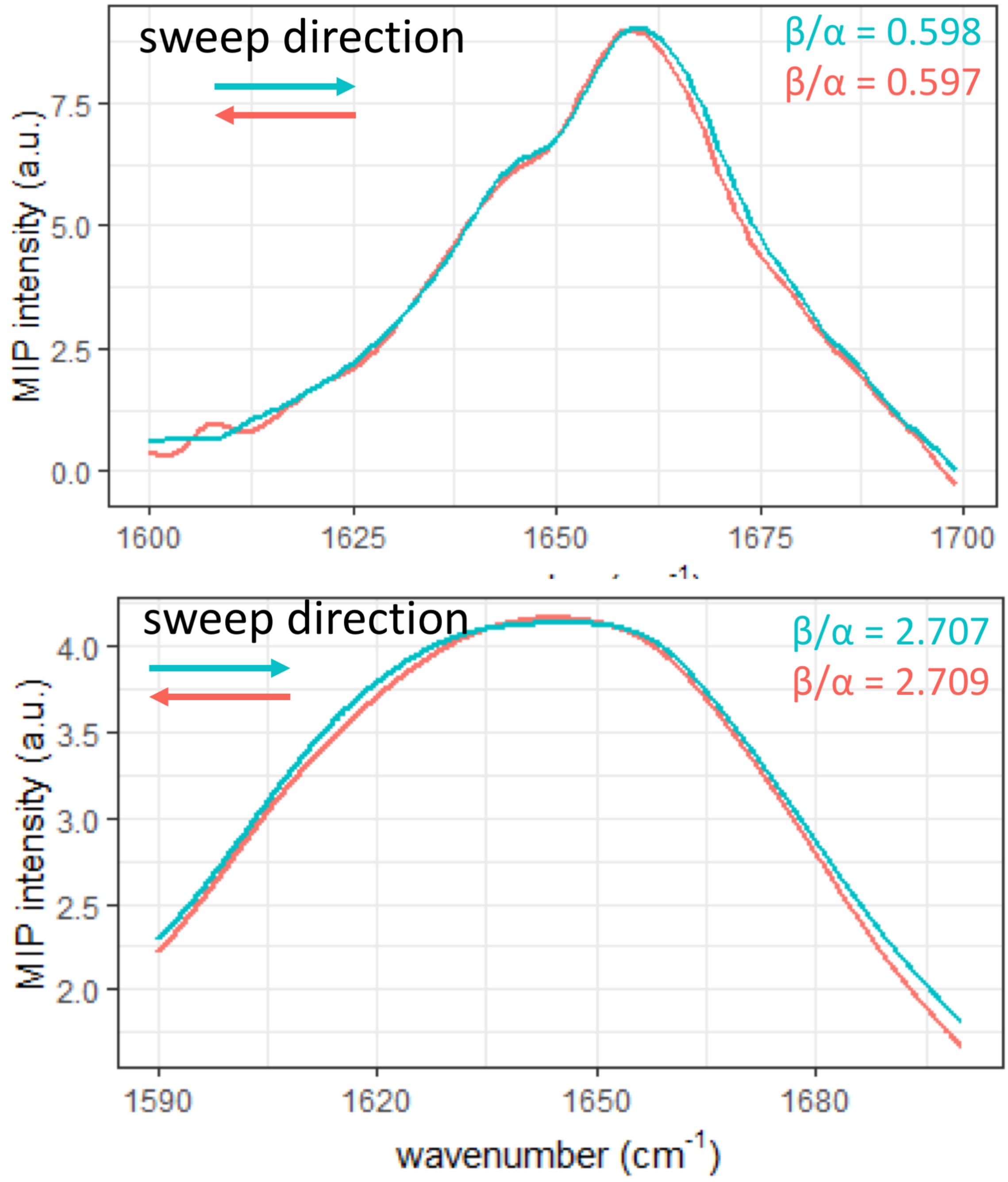


**Fig. S4.** Sweep wavenumbers from both directions did not affect MIP spectrum and β/α ratio quantification. MIP spectra acquired by sweeping both directions for (A) bovine serum albumin (BSA) powder and (B) htt103Q-Crimson aggregates in live yeast cells. Sweeping direction did not affect the MIP spectrum profile or the quantification of β/α ratio. It suggested that heat accumulation between mid-IR pulses should not be a concern.

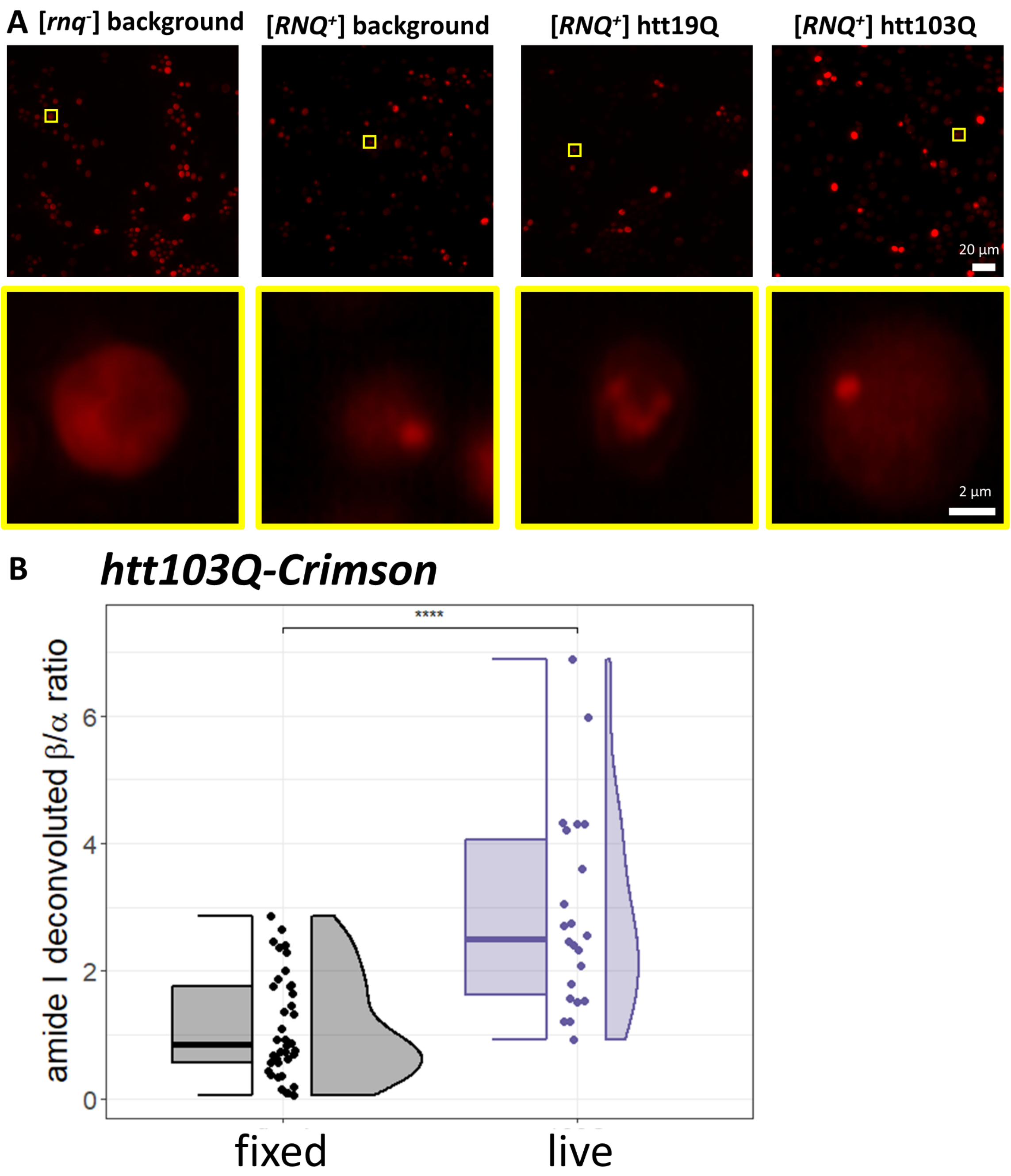


**Fig. S5.** PROTEOSTAT staining cross-validated aggregation. However, fixation affected β-sheet detection by MIP. (A) Representative fluorescence images for backgrounds [*rnq*$^-$] and [*RNQ*$^+$], as well as htt19Q and htt103Q in [*RNQ*$^+$]. Top: large field of view images. Bottom: details from yellow boxes. PROTEOSTAT showed diffuse fluorescence in prion free strain while strong fluorescent foci were observed for htt103Q. [*RNQ*$^+$] prion were also stained in non-transformed strains (wt). (B) β/ɑ ratio for fixed and live htt103Q-Crimson aggregates. Fixation caused low β/ɑ ratio measured by MIP. It is possible that to the cross-link of protein by paraformaldehyde affected secondary structure detection by MIP (1, 2).

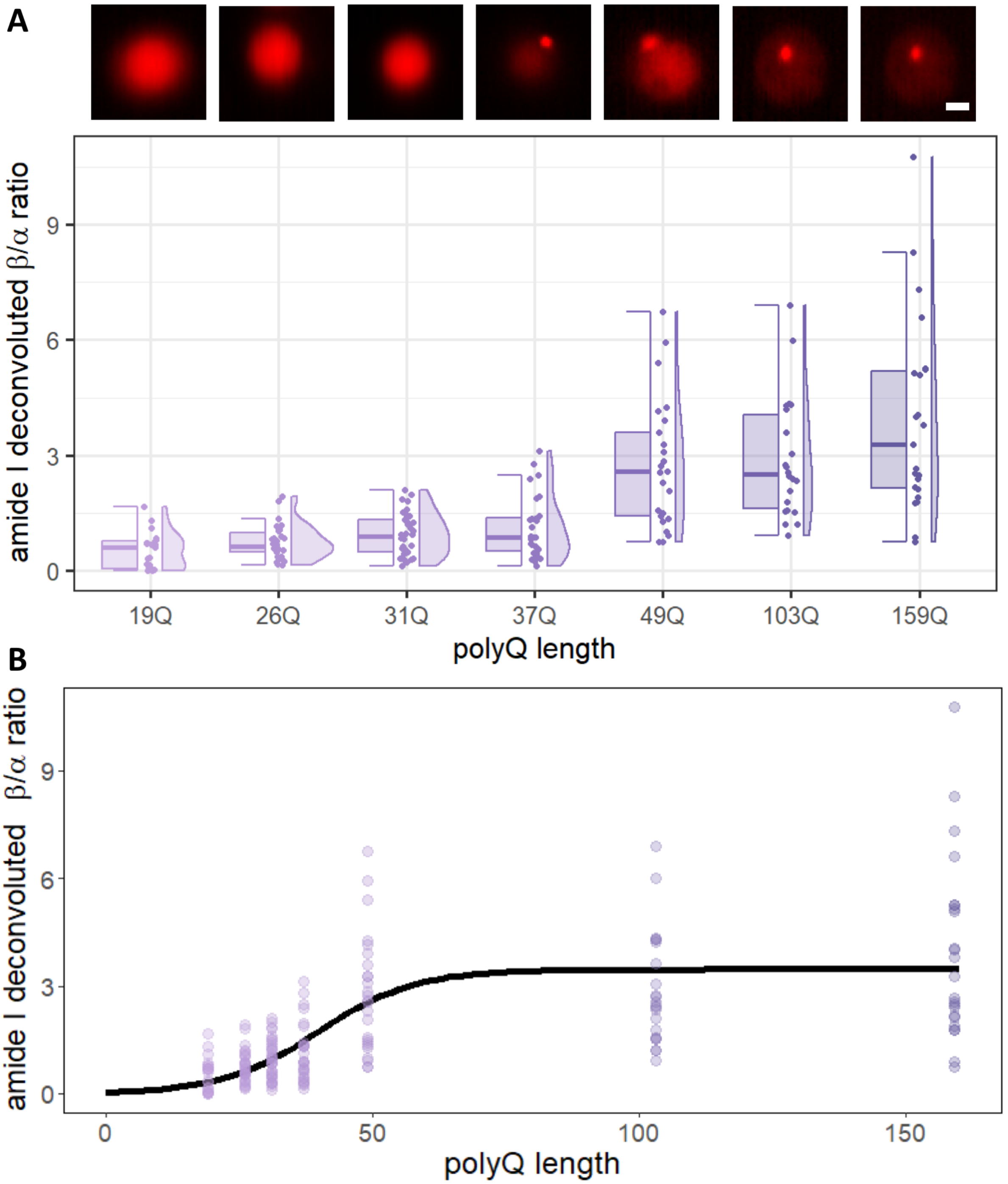


**Fig. S6.** β/α ratios for httnQ sweeping from low to high polyQ lengths. (A) β/α ratios for 19Q, 26Q, 31Q, 37Q, 49Q, 103Q and 159Q under Crimson fluorescent guidance. Notably, 37Q is just below the established threshold for toxicity of 38Q repeats. Top row shows representative Crimson fluorescent images for yeast cells expressing httnQ-Cr. Note that Crimson fluorescent foci started to appear at 37Q. (B) Sigmoidal curve (in black) correlating polyQ length with β/α ratios showed a graduate increase then consistent for high polyQ aggregates.

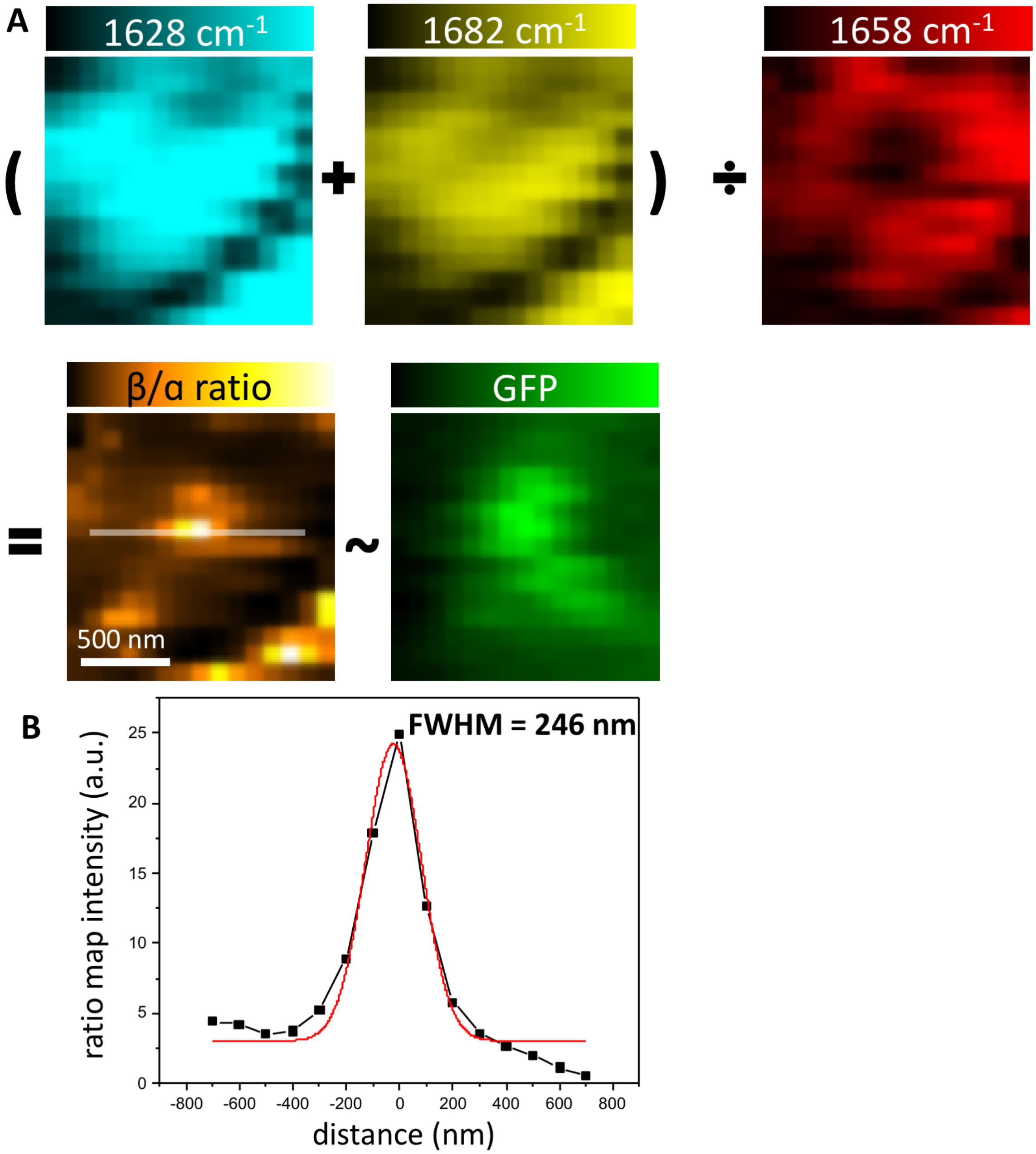


**Fig. S7.** Ratio map between MIP images revealed β-sheet enriched core below conventional optical diffraction limit. (A) ratio map for detail images in **Fig. 5*A***. The sum of low-wn value and high-wn value for β-sheet (1628 $cm^{-1}$ in cyan and 1682 $cm^{-1}$ in yellow) was divided by wn value for α-helix (1658 $cm^{-1}$ in red). The produced β/α ratio map did correlate with the GFP fluorescence image. (B) Gaussian fitting of the line profile from the β/α ratio map (in white) showed a full width at half maximum (FWHM) of 246 nm, below the theoretical limit of 270 nm

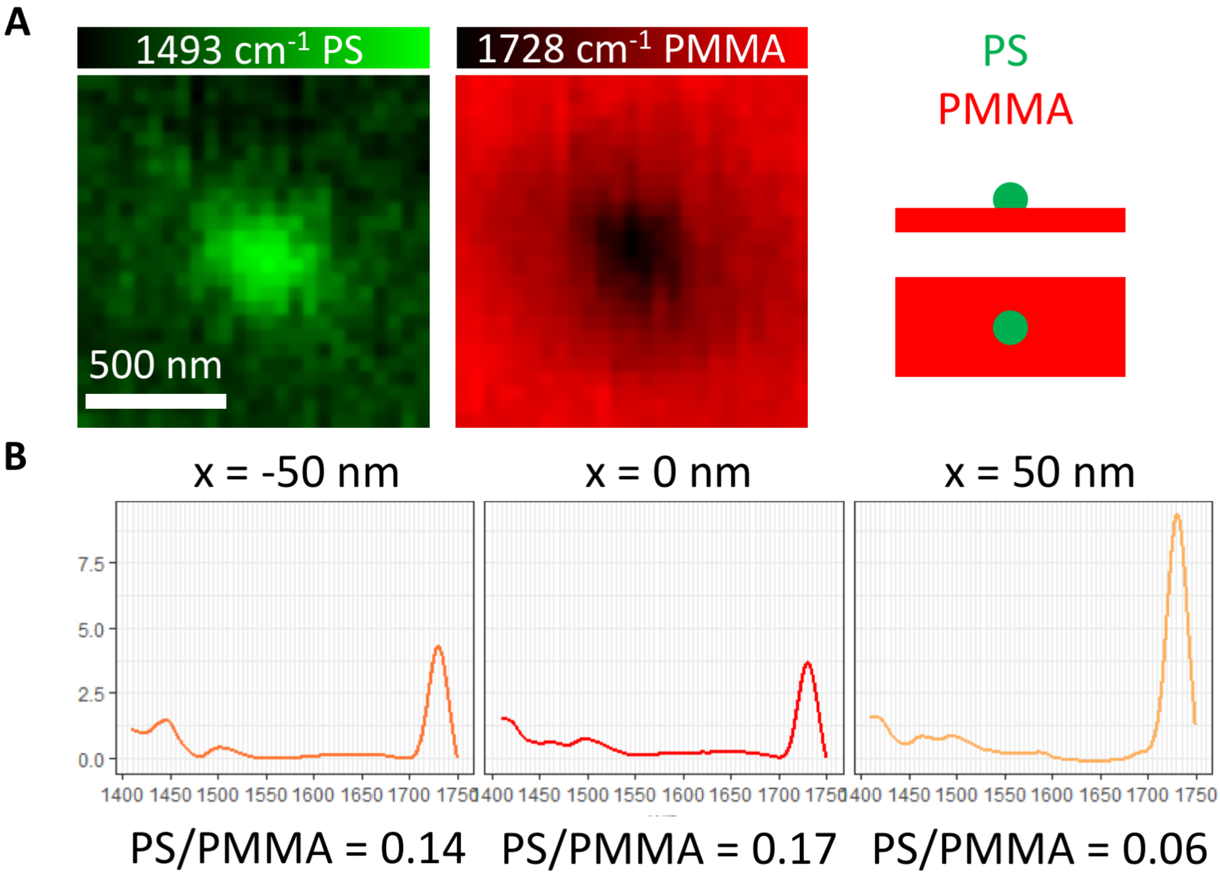


**Fig. S8.** To demonstrate the ability of MIP to distinguish spectral feature when two different chemical entities partition in a small core against a background, we recreated this scenario using a synthetic polymer mixtures: 200 nm in diameter polystyrene (PS) beads was deposited on top of spin coated Poly(methyl methacrylate) (PMMA) (A). Representative MIP image at 1493 cm$^{-1}$ (for PS) showed the single PS bead, while MIP image at 1728 cm$^{-1}$ (for PMMA) showed a corresponding void. (B) MIP spectra were acquired along the PS bead, with relative positions at -50 nm, 0 nm and +50 nm. The ratio of peak area integral at 1495 cm$^{-1}$ (for PS) divided by 1728 cm$^{-1}$ (for PMMA) after manual baseline correction were calculated as PS/PMMA ratio, to emulate β/ɑ ratio. PS/PMMA showed a similar trend of high intensity at PS wavenumber at the center of the bead, while decreasing intensity away from the center.

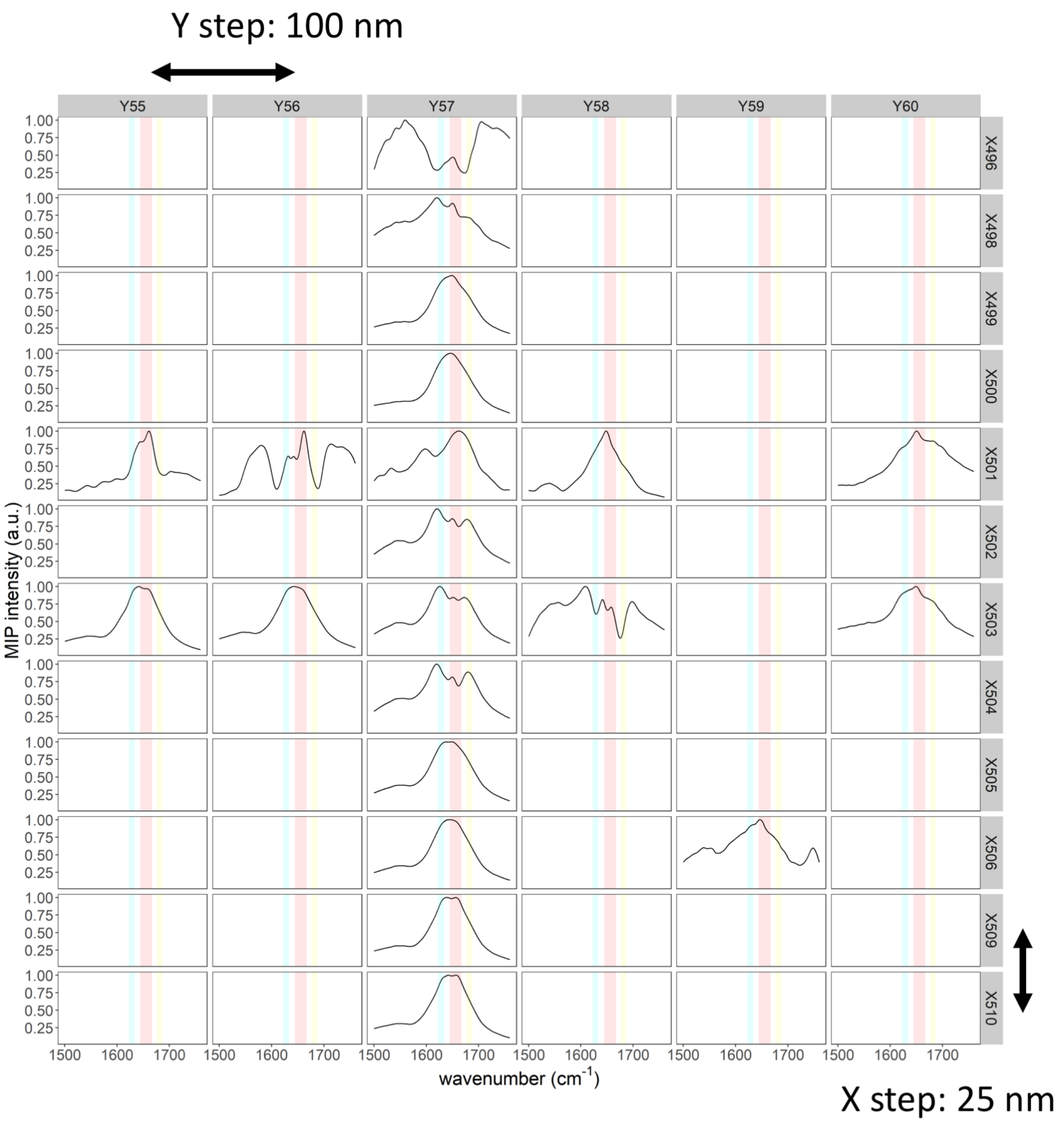


**Fig. S9.** An extended grid view of MIP spectra acquired with small spatial steps around one single aggregate of htt103Q-GFP. The Y step is 100 nm and the X step is 25 nm. Within each panel, the shaded lines represent low wn β-sheet (in cyan), α-helix (in red) and high wn β-sheet (in yellow). A clear continuous trend could be observed.

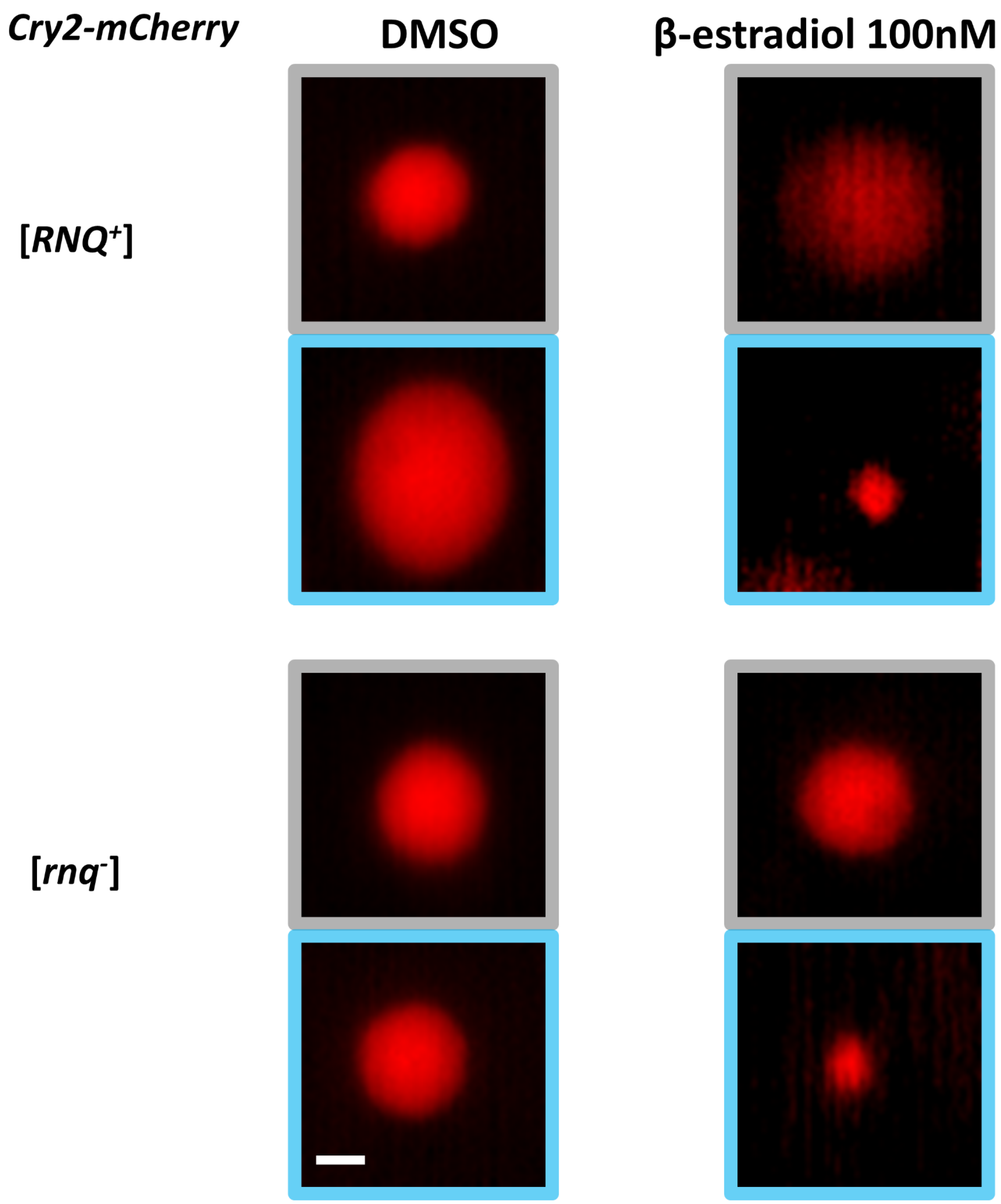


**Fig. S10**. Blue light stimulated oligomerization of Cry2-mCherry induced by β-estradiol. Representative mCherry fluorescence images (in red) were shown before (framed in gray) and after 5 min blue light stimulation (framed in blue). Only estradiol-induced cells show clear foci formation after blue light stimulation, while uninduced cells (DMSO control) show diffuse fluorescence. Scale bar: 2 μm.

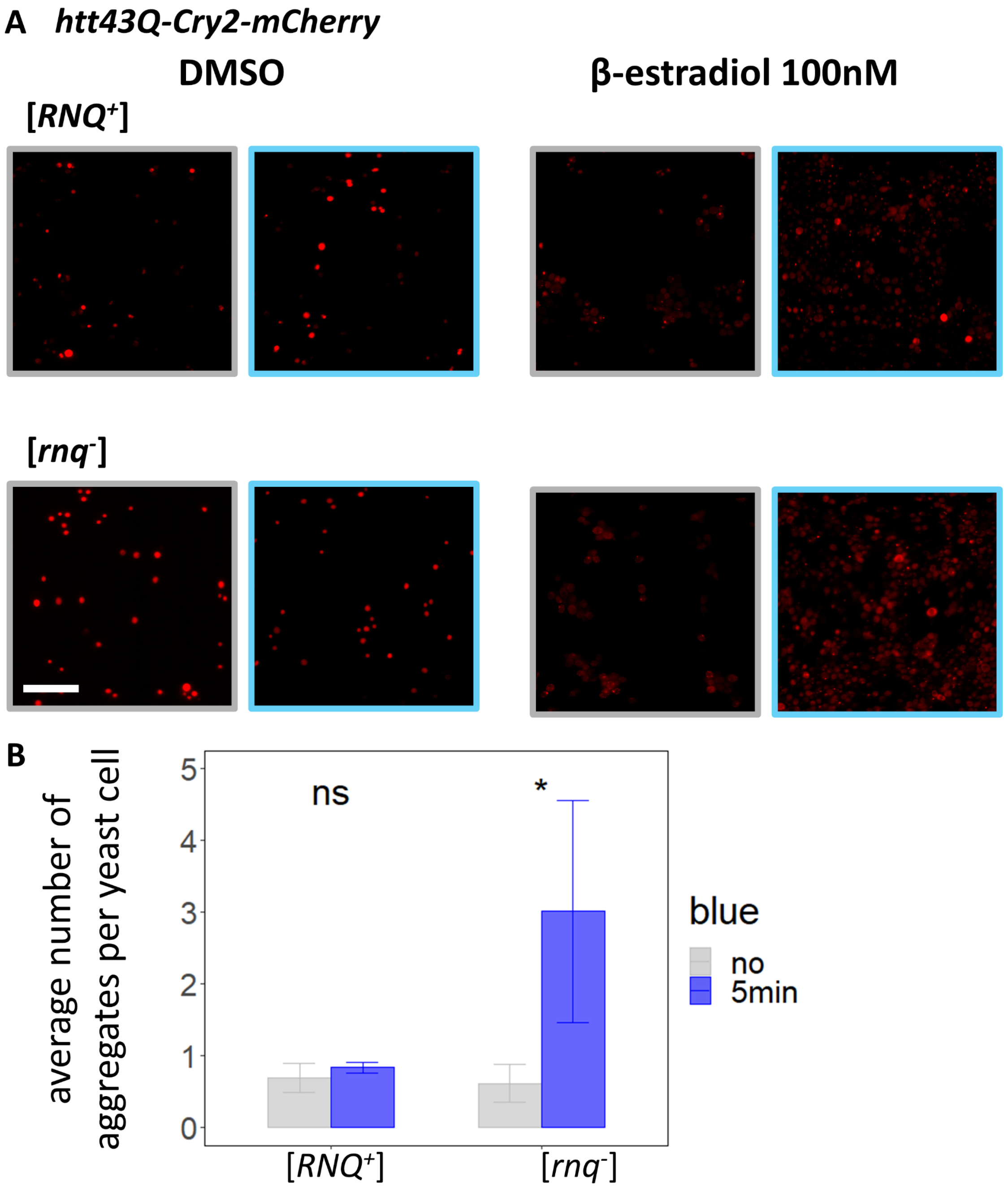


**Fig. S11.** Blue light further stimulated aggregation of htt43Q-Cry2-mCherry in β-estradiol induced [*RNQ*$^+$] and [*rnq*-] backgrounds. (A) Representative mCherry fluorescence images for htt43Q aggregates before (framed in gray) and after 5 min blue light stimulation (framed in blue). Scale bar: 50 μm. (B) Quantification of average number of aggregates per yeast cell, obtained by dividing the number of fluorescent foci by number of yeast cells. Blue light stimulation increased the frequency of aggregation in both β-estradiol induced strains with a significant difference in [*rnq*-] strain.

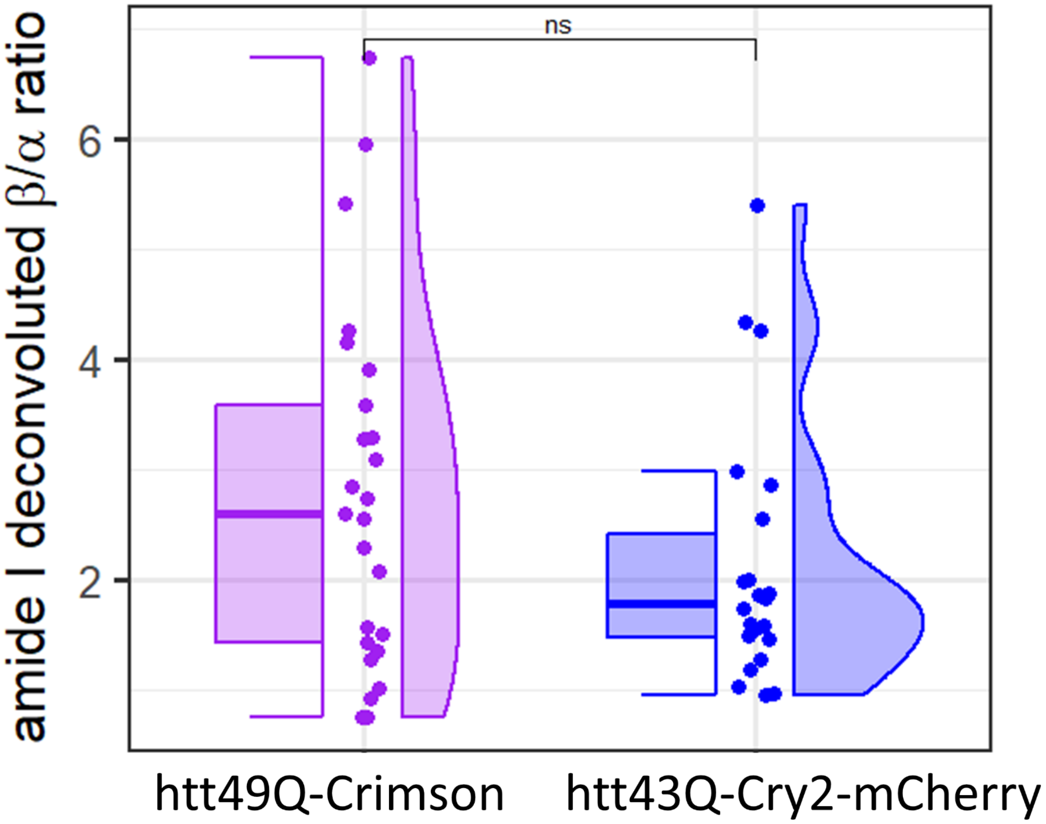


**Fig. S12.** β/α ratios for similar polyQ length without and with optogenetic modification. No significant difference between htt49Q-Crimson (N = 25) and htt43Q-Cry2-mCherry (N = 22) is observed. Unpaired pairwise t-test, $p = 0.122$, ns.

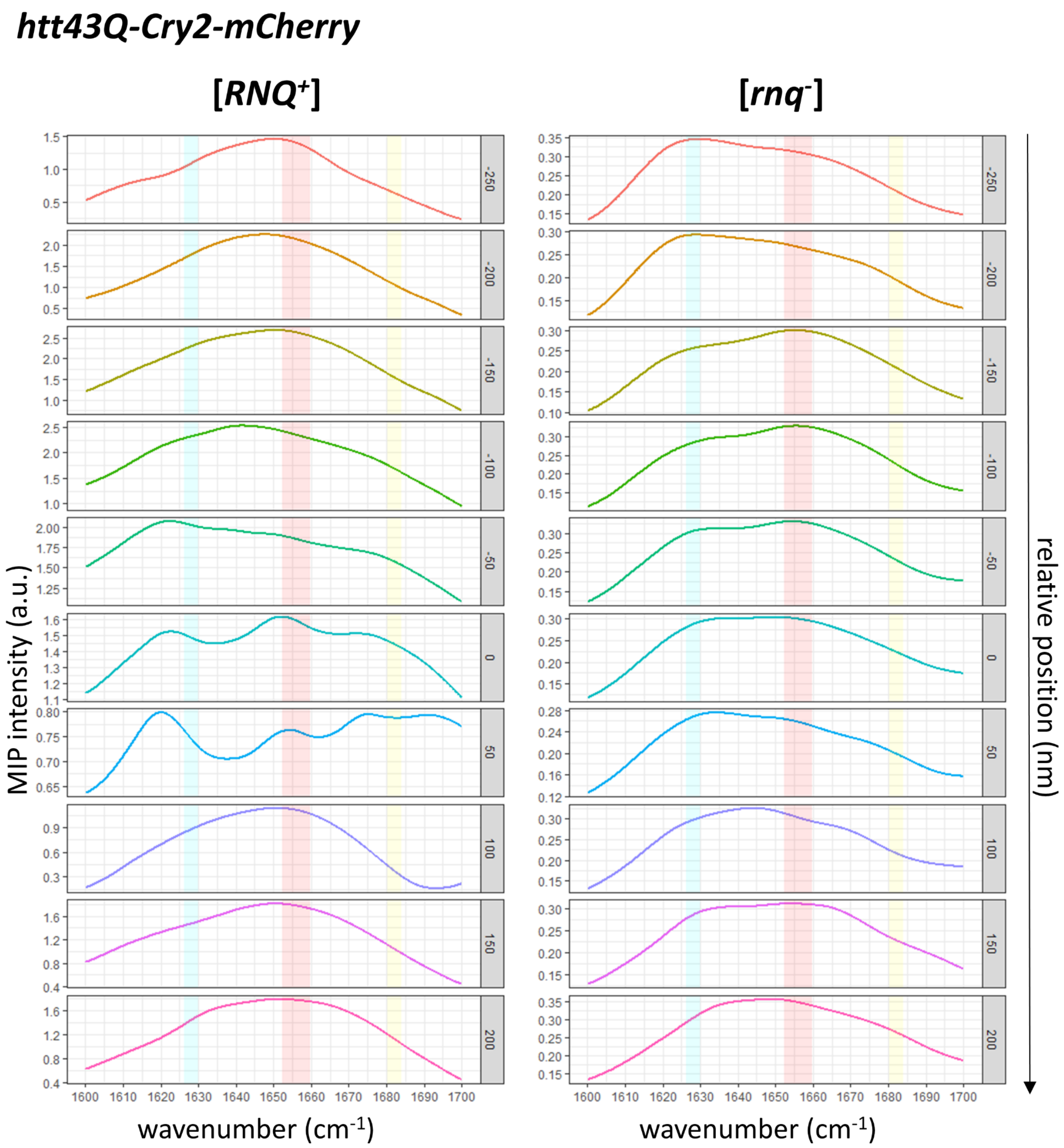


**Fig. S13.** MIP spectra in the amide I region of htt43Q-Cry2-mCherry aggregates in [*RNQ*$^{+}$] (left) and [*rnq*$^{-}$] (right) backgrounds. The MIP spectra were acquired across the aggregates with 50 nm step and β/α ratios were quantified across the aggregates shown in **Fig. 6*B***. Spectra from aggregates in [*RNQ*$^{+}$] strains showed significant β-sheet-related changes at the center and a gradual transition into broad amide I bands moving outwards. In contrast, MIP spectra from aggregates in [*rnq*$^{-}$] strains showed alternating spectral features with or without β-sheet related changes, suggesting multiple β-sheet core.

***htt43Q-Cry2-mCherry***

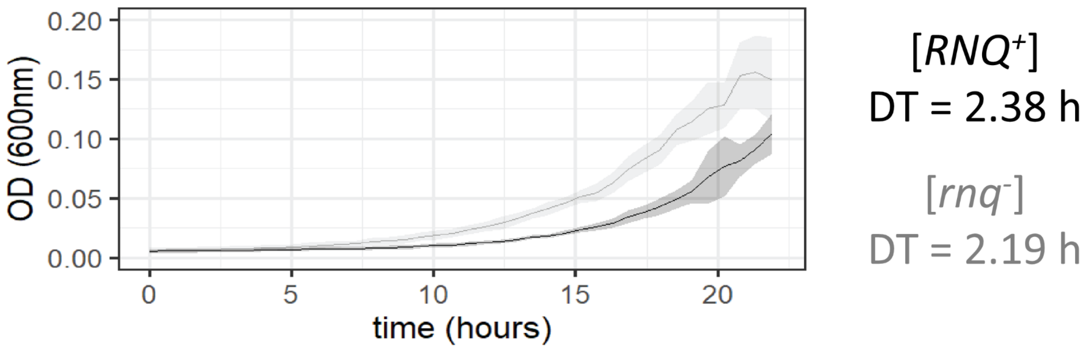


**Fig. S14.** Growth curve for yeast cells expressing htt43Q-cry2-mCherry with blue light stimulation in [*RNQ*$^+$] background (in black) and in [*rnq*$^-$] background (in gray). DT: doubling time. The solid line represents the means, and the shaded area represents the standard deviations for 6 replicate wells. Growth in [*RNQ*$^+$] was slower compared with in [[*rnq*$^-$] background.

#### Supplementary Methods

**PROTEOSTAT® Staining and fixation**. PROTEOSTAT® staining was performed using the ENZ-51023-KP050, Enzo kit. Briefly, yeast cells were first fixed with 4% paraformaldehyde in PBS for 15 min at room temperature. Then cells were permeabilized by 0.1% Triton X-100 in PBS for 20 min. Staining was performed by incubating treated cells for 15 min with 1:2000 dilution of PROTEOSTAT detection reagent, protected from light. Fluorescence imaging was performed with the same settings as for imaging httnQ-Cr protein fusion) (**Fig. S5*A***). htt103Q-Crimson cells were fixed with 4% paraformaldehyde in PBS for 15 min at room temperature (**Fig. S5*B***).

**Growth curve measurement with blue light and fitting.** The growth curve in **Fig. S14** for yeast cells expressing htt43Q-Cry2-mCherry in [$RNQ^+$] and [$rnq^-$] strains were measured in the 96-well plate by the microplate reader (SpectraMax I3x, Molecular Devices). To achieve blue light stimulation at the same time, the microplate reader was programmed to cycle through absorption measurement at 488 nm (shines blue light one column at a time) and shaking to facilitate yeast growth. The absorption of 600 nm light (OD 600 nm) was measured every 33.4 mins between cycles of blue light and shaking. All strains were grown from frozen glycerol stock overnight and diluted 1:500 in SD-ura with 100nM β-estradiol. 100 μl of diluted yeasts were added into each well of the 96-well plate with 6 replicates for each strain. The means and standard deviations were calculated for the 6 replicate wells at each time point (**Fig. S1** and **Fig. S14**). The doubling time were estimated from the averaged growth curve using R package GrowthCurver (3).
